# Gentamicin B1 is not a minor gentamicin component with major nonsense mutation suppression activity

**DOI:** 10.1101/476564

**Authors:** Alireza Baradaran-Heravi, David E. Williams, David A. Powell, Aruna D. Balgi, Raymond J. Andersen, Michel Roberge

## Abstract

Nonsense mutations are single base substitutions that introduce a premature termination codon (PTC) preventing the formation of full-length protein. They are the causative mutations in about 10% of patients in a large number of rare genetic diseases. High concentrations of the antibiotic gentamicin can induce the incorporation of an amino acid at a PTC and formation of full-length protein, a process called PTC readthrough. Gentamicin is composed of several related aminoglycosides. We recently reported (<u>doi/10.1073/pnas.1620982114</u>) that the major gentamicin components that are responsible for its antibacterial activity showed weak to no PTC readthrough activity but that the minor component gentamicin B1 was a potent readthrough inducer. We have now determined that gentamicin B1 acquired from the sole supplier at the time the study was carried out was not gentamicin B1 but instead the closely related aminoglycoside G418. Gentamicin B1 recently became available from a second commercial source. Here, we provide nuclear magnetic resonance (NMR) assignment data for the two commercial compounds and verify only the second is indeed gentamicin B1. We show that gentamicin B1 lacks PTC readthrough activity in HDQ-P1 and DMS-114 cells homozygous for the *TP53* R213X nonsense mutation, as well as in a cell-free translation assay.

## Introduction

Nonsense mutations in the coding sequence of a gene change an amino acid codon to a premature termination codon (PTC) and are responsible for about 10% of rare genetic disease cases (1). Compounds that permit insertion of an amino acid at the PTC enable formation of full-length protein and increased protein function, instead of truncated inactive protein. This strategy, termed nonsense suppression or PTC readthrough, has the potential to treat large numbers of patients across multiple rare genetic disorders. However, drugs that can induce therapeutic levels of readthrough at safe doses are not yet available.

Gentamicin is an aminoglycoside antibiotic approved for use in humans. It can induce PTC readthrough in cultured cells but only at concentrations that are orders of magnitude higher than the plasma concentration considered safe in humans (2). Clinical trials in Duchenne muscular dystrophy and cystic fibrosis patients have indicated that therapeutically relevant levels of readthrough cannot be achieved at subtoxic gentamicin doses (3–5). Gentamicin is not a pure compound but a mixture of related aminoglycosides isolated from the soil fungus *Micromonospora purpurea*. It is composed of specified proportions of the major gentamicins C1, C1a, C2, C2a and C2b. The U.S. Pharmacopeial Convention (USP) requires 25-50% C1; 10-35% C1a; 25-55% C2 + C2a while the European Pharmacopea specifies 20-40% C1, 10-30% C1a, 40-60% C2 + C2a + C2b (6). Gentamicin may also contain small amounts of a number of related aminoglycosides found in variable proportions in different drug batches and accounting together for < 3% of the material (6). Variable response to gentamicin treatment has been observed in animal models and humans (5, 7) and it has been speculated that this variability may be due to differences in gentamicin composition (7).

In a 2017 publication, we examined this question by measuring the activity of pure samples of each individual major gentamicin component and of minor gentamicin components (8). We found that none of the major gentamicins showed significant activity, an observation recently corroborated by others (9). We reported that the minor component gentamicin B1 was 30-to 100- fold more potent than the most potent gentamicin batches tested (8). In recent structure-activity relationship studies to be reported elsewhere, we observed that synthetic aminoglycoside analogs bearing ring 1 of gentamicin B1 lacked readthrough activity whereas analogs bearing ring 1 of the closely related aminoglycoside G418 showed readthrough activity. This puzzling observation led us to reexamine the structure and PTC readthrough activity of gentamicin B1.

## Materials and Methods

### NMR

All NMR spectra were recorded on a Bruker AV-600 spectrometer with a 5 mm CPTCI cryoprobe using standard Bruker pulse sequences. ^1^H chemical shifts are referenced to the residual DMSO-*d*_6_ (δ 2.49 ppm) and ^13^C chemical shifts are referenced to the DMSO-*d*_6_ solvent peak (δ 39.5 ppm).

### Human cells

The HDQ-P1 cell line was purchased from the German Collection of Microorganisms and Cell Cultures (DSMZ). DMS-114 cells were purchased from the American Type Culture Collection (ATCC). Both have homozygous nonsense mutation in the *TP53* gene (NM_000546.5:c.637C > T; NP_000537.3:p.R213X). HDQ-P1 and DMS-114 cells were cultured in high glucose Dulbecco’s modified Eagle medium (DMEM, Sigma-Aldrich) and RPMI-1640 (Sigma-Aldrich), respectively, supplemented with 10% (vol/vol) FBS (Sigma-Aldrich) and 1× antibiotic–antimycotic (Gibco/Thermo Fisher Scientific) at 37 °C and 5% (vol/vol) CO_2_.

### Automated Capillary Electrophoresis Western Analysis

Detection of p53 was performed as previously described (10). Briefly, HDQ-P1 or DMS-114 cells were seeded in 12-well culture plates and incubated with various concentrations of compounds. After 48-72 h incubation the cells were washed with ice-cold PBS and lysed in 50 µl lysis buffer composed of 20 mM Tris⋅HCl, pH 7.5, 150 mM NaCl, 1 mM EDTA, 1 mM EGTA, 1% (vol/vol) Triton X-100, 2.5 mM sodium pyrophosphate, 1 mM β-glycerophosphate, 1 mM Na_3_VO_4_, 1 mM DTT, and 1× complete protease inhibitor mixture (Roche Molecular Biochemicals). Lysates were centrifuged at 18,000 g for 15 min at 4 °C and protein level was quantitated in the supernatants using the Bradford assay and adjusted to 0.9 or 0.7 mg/ml protein for HDQ-P1 and DMS-114 cells, respectively. Capillary electrophoresis western analysis was carried out following the manufacturer’s instructions (ProteinSimple WES) using DO-1 p53 antibody from Santa Cruz Biotechnology and vinculin antibody (clone 728526) from R&D Systems. The data were analyzed with the inbuilt Compass software (ProteinSimple).

### Aminoglycosides

G418 disulfate was purchased from Sigma (A1720). Gentamicin B1 (B1-MCC) isolated from gentamicin sulphate complex C was purchased from MicroCombiChem, Germany as free base and as sulfate salt. Synthetic gentamicin B1 acetate (B1-TRC) was purchased from Toronto Research Chemicals.

### In Vitro Transcription and Translation Assays

In vitro transcription and translation assays were performed as previously described (10) with minor modifications. In brief, pcDNA-6.2/V5-DEST vector expressing p53 R213X-TGA was linearized with FastDigest MssI restriction enzyme (Thermo Fisher Scientific) and subjected to RNA synthesis using the mMESSAGE mMACHINE T7 ULTRA Transcription Kit (Thermo Fisher Scientific). 500 ng of the 5′ capped and poly(A)-tailed RNA was subjected to in vitro translation using the One-Step Human Coupled IVT Kit (Thermo Fisher Scientific). This assay was carried out at 30 °C for 30 min in the presence or absence of various concentrations of gentamicin B1 or G418 according to the manufacturer’s instructions. Samples were diluted 10 times in water and 5 µl of each sample was subjected to automated capillary electrophoresis western analysis for p53 detection.

## Results

The commercial gentamicin B1 samples used in our 2017 study (8), here termed B1-MCC, were purchased as free base and as sulfate salt. They were provided with Certificates of Analysis, mass spectrometry and NMR data (Supplementary Figure 1) supporting designation as gentamicin B1, which was corroborated by an independent NMR analysis carried out at our request. During the summer of 2018, while carrying out structure-activity relationship studies of synthetic pseudotrisaccharides, we noticed that replacing ring 1 of G418 with ring 1 of gentamicin B1 caused loss of PTC readthrough activity (unpublished). This observation made us suspect the identity of B1-MCC.

**Figure 1.**
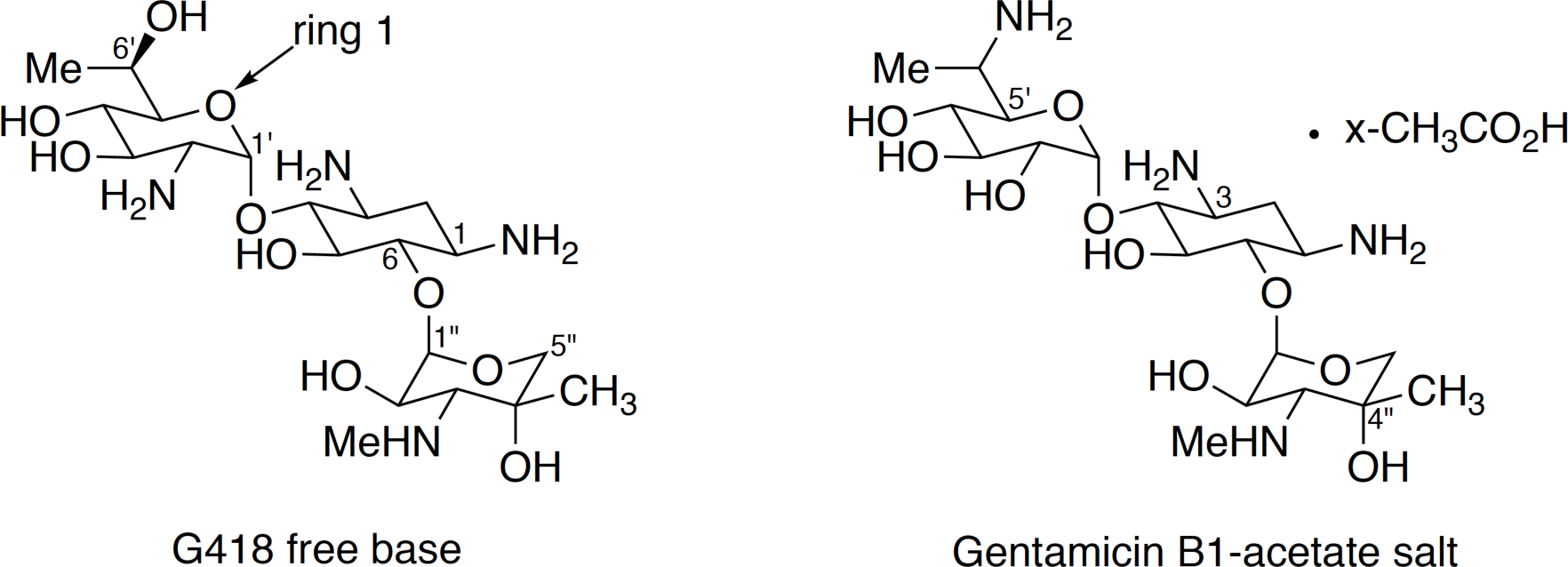
Structure of G418 and gentamicin B1 assigned from NMR data in Table 1.

Gentamicin B1 and G418 are closely related compounds with the same mass and molecular formula (496.55 g/mol; C_20_H_40_N_4_O_10_) that differ only in the positions of the NH_2_ and OH functional groups at C-2′ and C-6′ as shown in the structural drawings in Figure 1. We have been unable to find any published reference NMR assignments for gentamicin B1 in the peer-reviewed or patent literature. The Certificate of Analysis provided by the vendor included ^1^H and ^13^C NMR assignments for spectra recorded in D_2_O (Supplementary Figure 1). A resonance at δ 54.7 was assigned to a secondary alcohol carbon C-2′ and a resonance at δ 65.6 to an amine-bearing carbon C-6′. Support for the placement of the alcohol at C-2′ and the amine at C-6′ in this sample came from an ^1^H-^15^N-HMBC spectrum recorded in D_2_O at the Centre for Drug Research and Development in Vancouver. This spectrum showed a strong ^1^H/^15^N correlation that had a ^1^H chemical shift of δ 1.13 ppm. A methyl doublet at δ 1.13 in the 1D ^1^H NMR spectrum of this B1-MCC sample could be confidently assigned to C-7′ since it was the only methyl group in the molecule vicinal to a methine. They concluded that the strong ^1^H-^15^N-HMBC correlation between Me-7′ and a nitrogen was only possible if there was an amino functionality at C-6′, suggesting that the sample was gentamicin B1.

**Table 1.** ^13^C and ^1^H NMR data comparison for G418 free base, purchased as gentamicin B1 (B1-MCC) and gentamicin B1-acetate salt (B1-TRC) recorded in DMSO-*d_6_* and 3:2 DMSO-*d_6_*/D_2_O, respectively, at 600 MHz.

|  | **G418 free base (B1-MCC)** |  | **Gentamicin B1-acetate salt (B1-TRC)** |  |
| --- | --- | --- | --- | --- |
| **Position** | $^{13}\text{C}$ ($\delta$) | $^1\text{H}$ ($\delta$, multiplicity (J Hz)) | $^{13}\text{C}$ ($\delta$) | $^1\text{H}$ ($\delta$, multiplicity (J Hz)) |
| **1** | 51.7 | 2.59, bt (9.2) | 51.5 | 2.96, m |
| **2<sub>ax</sub>** | 38.7 | 0.99<sup>a</sup> | 33.0 | 1.33, m |
| **2<sub>eq</sub>** |  | 1.74, m |  | 2.00, m |
| **3** | 50.3 | 2.51<sup>a</sup> | 49.0 | 3.02, m |
| **4** | 92.2 | 2.89, t (9.2) | 83.0 | 3.37<sup>a</sup> |
| **5** | 74.2 | 3.34, t (9.2) | 73.9 | 3.49<sup>a</sup> |
| **6** | 86.1 | 2.97, t (9.2) | 86.5 | 3.26, t (9.1) |
| **1'** | 102.3 | 4.85, d (3.1) | 97.5 | 5.28, d (3.7) |
| **2'** | 56.2 | 2.52<sup>a</sup> | 72.4 | 3.36, dd (9.9, 3.7) |
| **3'** | 74.0 | 3.24, t (9.2) | 73.9 | 3.49<sup>a</sup> |
| **4'** | 72.9 | 3.01<sup>a</sup> | 71.8 | 3.82<sup>a</sup> |
| **5'** | 75.1 | 3.52, bd (7.8) | 70.7 | 3.16, t (9.6) |
| **6'** | 66.2 | 3.87<sup>a</sup> | 47.8 | 3.51<sup>a</sup> |
| **7'-Me** | 17.4 | 1.01, d (6.3) | 12.7 | 1.12, d (7.0) |
| **1''** | 100.3 | 4.92, d (3.2) | 101.6 | 4.88, d (3.6) |
| **2''** | 69.7 | 3.43, dd (10.2, 3.2) | 67.9 | 3.93, dd (10.8, 3.6) |
| **3''** | 64.2 | 2.28, d (10.2) | 65.0 | 3.07, d (10.8) |
| **4''** | 71.2 | / | 70.9 | / |
| **5''<sub>ax</sub>** | 67.4 | 3.89, d (11.9) | 68.4 | 3.82<sup>a</sup> |
| **5''<sub>eq</sub>** |  | 3.05, d (11.9) |  | 3.21, d (12.5) |
| **3''-NH-Me** | 38.1 | 2.45, s | 36.2 | 2.66, s |
| **4''-Me** | 23.9 | 0.97, s | 22.7 | 1.13, s |

<sup>a</sup>Multiplicity not determined due to overlapping signals - chemical shifts determined from 2D NMR data.

In summer 2018, synthetic gentamicin B1, here termed B1-TRC, became available as the acetate salt from a second commercial vendor. The Certificate of Analysis and vendor NMR data are shown in Supplementary Figure 2. We have carried out detailed 1D and 2D NMR analysis of both the B1-MCC and B1-TRC samples and our ^1^H and ^13^C assignments for both samples are listed in Table 1. Of particular note are our assignments of δ 56.2 for C-2′ and of δ 66.2 for C-6′ in the ^13^C NMR spectrum of the B1-MCC sample recorded in DMSO-*d_6_*. These values are in good agreement with the assignments of the ^13^C shifts of these two carbons provided by the vendor for spectra recorded in D_2_O [δ 54.7 (C-2’); 65.6 (C-6’)] (Supplementary Figure 1) allowing for minor shifts due to different solvents. However, the C-2′ resonance was more shielded and the C-6′ was more deshielded than expected/calculated for the proposed gentamicin B1 structure that has an alcohol at C-2′ and an amine at C6′. This suggested that the structure assigned by the vendor was incorrect and that the positions of the amino and alcohol functionalities needed to be switched, so that B1-MCC was actually G418. We then mixed B1-MCC with authentic G418 (from Sigma) and recorded 1D ^1^H NMR data on the mixture. This experiment showed that the two compounds were identical.

**Figure 2.**
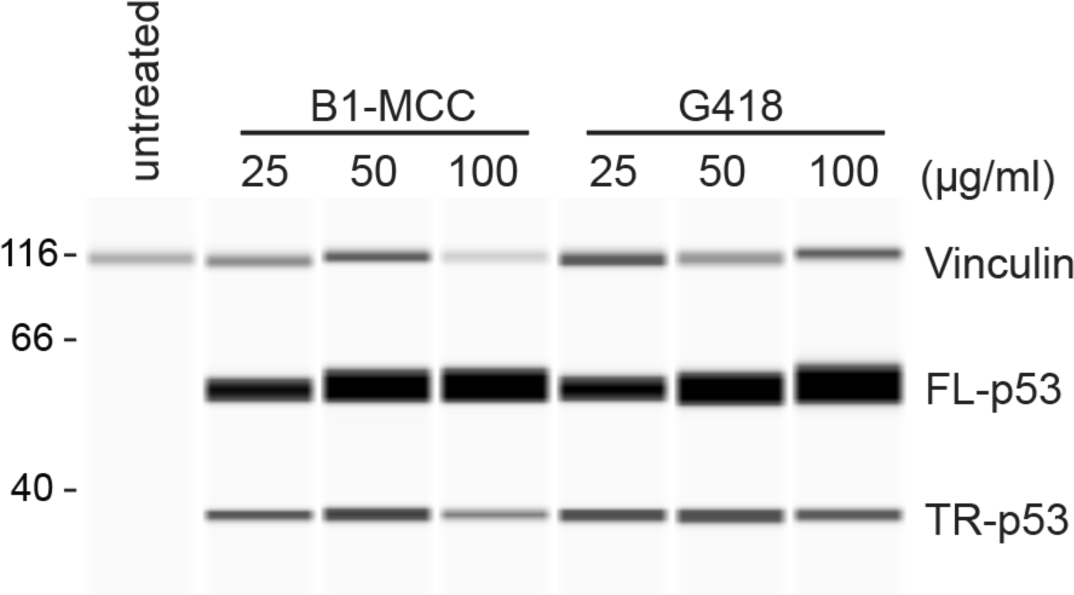
Comparison of the PTC readthrough activity of B1-MCC and G418. DMS-114 cells were incubated with the indicated concentrations of B1-MCC sulfate and G418 sulfate for 72 h and the production of full-length p53 (FL-p53) and truncated p53 (TR-p53) was determined using automated capillary electrophoresis western analysis with vinculin as a protein loading control. Molecular weight markers are shown on the left in kDa.

Our independent NMR structural analysis of the B1-TRC sample showed it is indeed gentamicin B1 (Table 1) and C-2′ bearing an alcohol was assigned to a carbon resonance at δ 72.4 and C-6′ now bearing an amino nitrogen was assigned to a carbon resonance at δ 47.8. Hence, the key to distinguishing between the two structures G418 and gentamicin B1 is the correct assignment of the carbon chemical shifts of C-6′ and C-2′ to carbons bearing either amino or alcohol functionalities. The configuration at C-6′ of B1-TRC remains undefined as it is not possible to assign this from the NMR analysis alone. Finally, it appears that the putative diagnostic ^1^H-^15^N-HMBC correlation observed at the Centre for Drug Research and Development arises from a correlation between H-2ax and one of the adjacent nitrogen atoms at either C-1 or C-3. The H-2_ax_ ^1^H resonance was presumably isochronous with the Me-7′ ^1^H resonance since we also observed nearly identical chemical shifts for Me-7′ (δ 1.01) and H-2_ax_ (δ 0.99) in our data for B1-MCC (Table 1).

We compared the PTC readthrough activity of B1-MCC and G418 from Sigma. DMS-114 cancer cells with homozygous R213X nonsense mutation in the *TP53* gene were analyzed for p53 expression using automated capillary electrophoresis western analysis. Untreated cells express no detectable full-length p53, the readthrough product (Fig. 2). Exposure to B1-MCC or G418 caused a similar concentration-dependent increase in full-length p53 (Fig. 2). This result is consistent with the assignment of B1-MCC as G418.

We then compared the activity of B1-TRC and G418. In DMS-114 cells, B1-TRC showed no PTC readthrough activity even at a high concentration of 200 µg/ml (Fig. 3A), whereas G418 did in the same experiment. Similarly, B1-TRC showed no PTC readthrough activity in HDQ-P1 cells, a second cancer cell line with homozygous R213X nonsense mutation in the *TP53* gene, while G418 did (Fig. 3B). Additionally, B1-TRC showed no PTC readthrough activity in a cell-free translation assay, whereas G418 did (Fig. 3C). These results indicate that gentamicin B1 has no readthrough activity.

**Figure 3.**
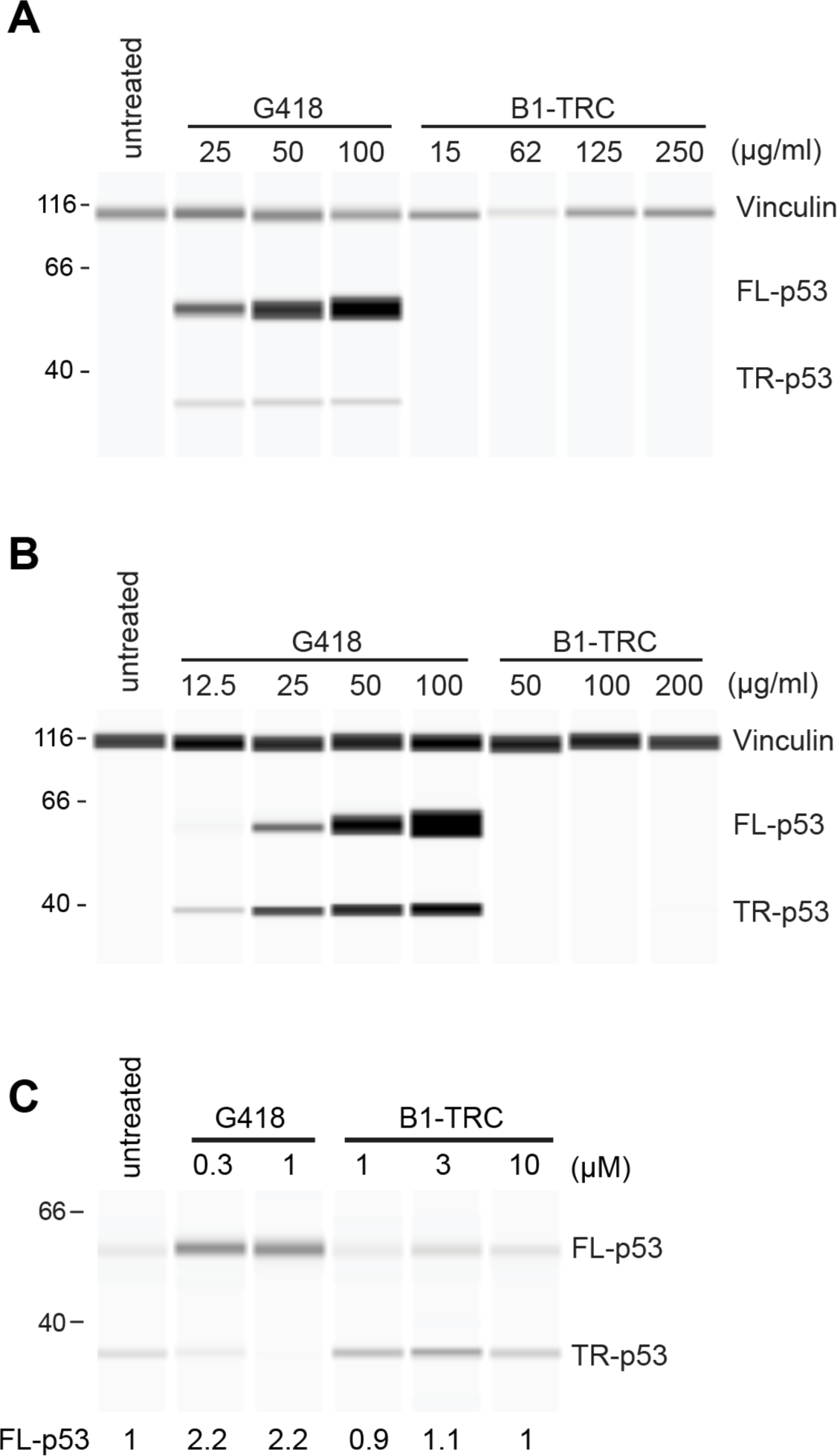
Effect of gentamicin B1 on PTC readthrough in cells and in a cell-free assay. DMS-114 cells (A) and HDQ-P1 cells (B) were incubated with the indicated concentrations of G418 sulfate or gentamicin B1 acetate for 48 h (A) or 72 h (B) and production of full-length p53 (FL-p53) and truncated p53 (TR-p53) was determined using automated capillary electrophoresis western analysis. Vinculin was used as a protein loading control. (C) Synthetic R213X *TP53* mRNA was subjected to in vitro translation in the presence of the indicated concentrations of G418 sulfate or gentamicin B1 acetate for 30 min and p53 was detected as in A and B. FL-p53 signal intensity relative to untreated control is shown below the lanes.

## Discussion

We previously reported that the major components of pharmaceutical gentamicin showed only weak PTC readthrough activity (8), corroborated recently by Friesen (9), but that the minor component B1was a potent readthrough compound (8). Here, we correct the record and show that the B1 used in our study was misidentified by the vendor and was in fact G418, and we provide evidence that synthetic gentamicin B1 is inactive as a readthrough compound.

Recent X-ray crystallography studies highlight a key role of the 6′-OH in ring 1 of G418 for binding to the ribosomal decoding center and eliciting a conformational change that enables PTC readthrough (11, 12). The authors speculate that replacing the 6′-OH group in ring 1 of G418 with a 6′-NH_2_ group, as occurs in B1, could preclude ribosomal binding (12). The proposed importance of the 6′-OH group for PTC readthrough is supported by the experimental data we present here comparing G418 with synthetic gentamicin B1, and by a recent study showing that introduction of a 6′-NH_2_ group in ring 1 of synthetic pseudotrisaccharides significantly reduces PTC readthrough activity (13).

## Supporting information

