## Supplementary material for "Gentamicin B1 is not a minor gentamicin component with major nonsense mutation suppression activity"

micro combichem
compound libraries for pharmaceutical research

**MicroCombiChem e.K.**
Rheingaustr. 190-196, E512
D-65203 Wiesbaden, Germany


### CERTIFICATE OF ANALYSIS

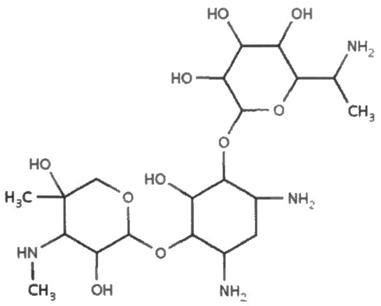

| | |
|---|---|
| **Product Name:** | **Gentamicin B1 Free Base** |
| **Article No.:** | **MCC3436** |
| Formula Weight: | 496.55g/mol |
| CAS: | [36889-16-4] |
| Formula: | C20H40N4O10 |
| Isotopic Mass$_{Calc.}$: | 496.2744 |
| Producer: | MicroCombiChem e.K. |
| Procedure: | Isolation from Gentamicin Sulphate Complex C |

| | |
|---|---|
| Chemical Name (IUPAC): | 2-(1-aminoethyl)-6-[(4,6-diamino-3-{[3,5-dihydroxy-5-methyl-4-(methylamino)oxan-2-yl]oxy}-2-hydroxycyclohexyl)oxy]oxane-3,4,5-triol |
| **Lot:** | **RegID 8070** |
| Production Time: | February/ March 2016 |
| Appearance: | White amorphous powder |
| Identity: | HPLC-MS (ESI); NMR; |
| Chromatographic Purity: | Min.98% (ELSD / MS) |
| MS: | Conforms |
| ELSD (Evaporative Light Scattering Detector): | Conforms |
| 1H-NMR: | Conforms |
| NMR-Purity: | Min.98% |
| Water Content: | 0.14% (Karl-Fischer) |
| Melting Point: | 68°C (instrument: Büchi 510) |
| Certified Purity: | Min.97% |
| Date of Analysis: | April 2016 |
| Storage (Short Term): | 0 to 4°C, dark, dry, inert gas; min. 2 weeks |
| Storage (Long Term): | -20°C, dark, dry, inert gas; min.5 years |
| Stability: | Estimated stability: minimum 5 years. |
| Recommended re-test procedures: | Quality control every third year by NMR / HPLC-MS. |
| Hazard Codes: | Not available yet. |

**COMMENTS:**

Packed under nitrogen. High stability at room temperature for min.14 days in solid and neutral solution form (ACN / H2O1:1), without any traces of decomposition. The compound has no significant UV-absorbance.

The estimation of the purity was made using HPLC-MS-ELSD (ESI+) and NMR.

For analytical testing use preferably Waters Atlantis T3 type columns or equivalent and min.pH2

For quality assurance purposes, for long term store the compound at min.-20°C

This product is intended for investigational laboratory use only. It is pharmaceutically unrefined and may contain traces of uncharacterized toxic impurities. It is not intended for use in humans or animals, not for food, drug or household use. Responsibility for its use and compliance with all applicable laws rests solely with the purchaser.

01.04.2016 A. Kl[illegible]

MicroCombiChem e.K.
- Compound Libraries -
Rheingaustr. 190-196, E512
D-65203 Wiesbaden


TRADE REGISTER: AMTSGERICHT WIESBADEN HRA 8724, GERMANY ♦ TAX ID 040 836 00249 ♦ VAT ID DE253555677 ♦ MANAGEMENT: DR. ANETTE KLINGER
BANK DETAILS: NASSAUISCHE SPARKASSE WIESBADEN ♦ CODE 510 500 15 ♦ ACC 135 215 374 ♦ S.W.I.F.T. NASS DE 55 ♦ IBAN DE80 5105 0015 0135 2153 74

### NMR Analysis

#### Sample BatchID: MCC3436 ‘6.8g-Charge’, Gentamicin B1 Disulfate 33.7%
#### Date: 26-02-2018
#### Analysis Number: 2018 - 11

Spectra at Bruker Fourier 400
Solvent: D2O
(1/ $^{1}$H)
(2/ $^{13}$C)
(3/ cosy)
(4/ hsqced)
(5/ hmbc)
(6/ dept)

#### Evidence of Chemical Structure and Assignment of NMR Spectra

<u>Results for Batch MCC3436 ‘6.8g-Charge’ Gentamicin B1 Disulfate 33.7%:</u>

The NMR analytical data of batch MCC3436 ‘6.8g-Charge’ are conform with the structure of Gentamicin B1.
$^{1}$H and $^{13}$C chemical shifts have been assigned using correlation spectra (cosy, hsqc, hmbc, dept). There are no significant conflicting correlations observable in the spectra. Traces of gentamicin congeners, acetonitrile and formic acid are present.

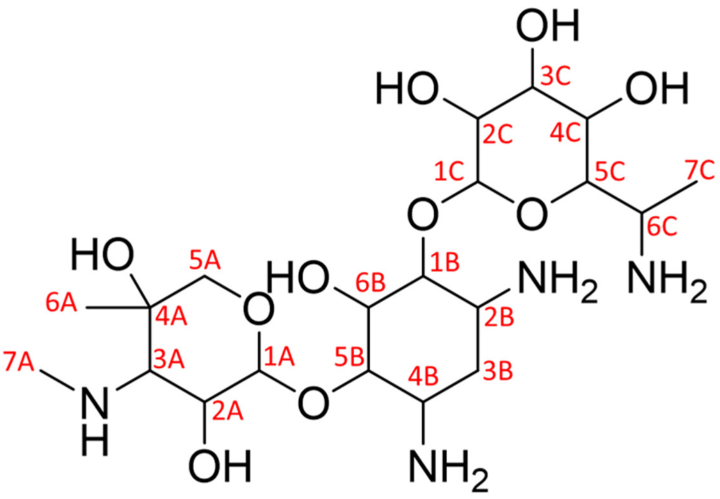

*Figure 1: Structure scheme of Gentamicin B1 and atom numbering used for this report.*

*Table 1: Shifts and proposed assignment. The shift values are not absolute.*

| | Pos. | δ ($^{13}C$) [ppm] | m($^{13}C$) | δ ($^{1}H$) [ppm] | mult | $J_{HH}$ coupling constant and partner | $^{n}J_{CH}$ correlations; δ ($^{13}C$) and assignment |
|---|---|---|---|---|---|---|---|
| A | 1 | 100.4 | CH | 5.05 | d | 3.8 Hz (A2) | 5.05 ($^{1}J$, 173 Hz, A1), 3.98 ($^{2}J$, $A5_{eq}$), 3.37 ($^{2}J$, $A5_{ax}$), 3.27 ($^{2}J$, A3) |
| | 2 | 66.6 | CH | 4.07 | dd | 10.9, 3 (A3), 3.8 Hz (A1) | 3.27 ($^{2}J$, A3), 5.05 ($^{2}J$, A1) |
| | 3 | 63.7 | CH | 3.27 (3.33-3.23) | m | 10.8 Hz (A2) | 5.05 ($^{3}J$, A1), 4.07 ($^{2}J$, A2), 3.37 ($^{3}J$, $A5_{ax}$), 2.80 ($^{4}J$, A7) |
| | 4 | 70.1 | C | - | - | - | 3.98 ($^{2}J$, $A5_{eq}$), 3.37 ($^{2}J$, $A5_{ax}$), 1.26 ($^{3}J$, A6) |
| | 5 | 67.4 | $CH_2$ | $3.98_{eq}$<br>$3.37_{ax}$ | d<br>d | 12.9 Hz ($A5_{ax}$)<br>12.8 Hz ($A5_{eq}$) | 5.05 ($^{3}J$, A1), 1.26 ($^{2}J$, A6) |
| | 6 | 21.0 | $CH_3$ | 1.26 | s | - | 1.25 ($^{1}J$, 127 Hz, A7) 3.37 ($^{3}J$, $A5_{ax}$), 3.27 ($^{3}J$, A3) |
| | 7 | 35.0 | $CH_3$ | 2.80 | s | - | 2.80 ($^{1}J$, 143 Hz, A7), 3.27 ($^{3}J$, A3), 1.26 ($^{2}J$, A6) |
| B | 1 | 84.3 | CH | 3.47 | q | 9.4 Hz (B2, B6) | 5.33 ($^{2}J$, C1), 3.64 ($^{2}J$, B6), 2.17 ($^{3}J$, $B3_{eq}$), 1.49 ($^{3}J$, $B3_{ax}$) |
| | 2 | 50.2 | CH | 3.16 (3.20 – 3.06) | m | (B1,B3) | 3.47 ($^{2}J$, B1), 2.17 ($^{2}J$, $C3_{eq}$), 1.51 ($^{2}J$, $C3_{ax}$) |
| | 3 | 31.7 | $CH_2$ | $2.17_{eq}$<br>$1.49_{ax}$ | dt<br>dd | 12.8 ($B3_{ax}$), 4.3 Hz (B2, B4)<br>12.6 Hz ($B3_{eq}$, B2, B4) | 3.16 ($^{2}J$, B2), 3.07 ($^{2}J$, B4) |
| | 4 | 49.1 | CH | 3.07 (3.20 – 3.06) | m | (B3, B5) | 3.48 ($^{2}J$, B5), 2.17 ($^{2}J$, $C3_{eq}$), 1.51 ($^{2}J$, $C3_{ax}$) |
| | 5 | 84.9 | CH | 3.48 | q | 9.4 Hz (B4, B6) | 5.05 ($^{2}J$, A1), 3.64 ($^{2}J$, B6), 3.47 ($^{3}J$, B5), 2.17 ($^{3}J$, $B3_{eq}$), 1.49 ($^{3}J$, $B3_{ax}$) |
| | 6 | 73.8 | CH | 3.64 (3.59-3.69) | m | (B1, B5) | 3.48 ($^{2}J$, B5), 3.47 ($^{2}J$, B1) |
| C | 1 | 98.8 | CH | 5.33 | d | 3.9 Hz (C1) | 5.33 ($^{1}J$, 171 Hz, C1), 3.47 ($^{3}J$, B1), 3.02 ($^{2}J$, C2) |
| | 2 | 54.7 | CH | 3.02 | dd | 10.5 Hz (C3),3.9 Hz (C1) | 5.33 ($^{2}J$, C1), 3.63 ($^{2}J$, C3) |
| | 3 | 71.7 | CH | 3.63 | m | (C2, C4) | 5.33 ($^{3}J$, C1), 3.30 ($^{2}J$, C4), 3.02 ($^{2}J$, C2) |
| | 4 | 70.6 | CH | 3.30 (3.33-3.23) | m | (C3, C5) | 3.81 ($^{2}J$, C5), 3.63 ($^{2}J$, C3) |
| | 5 | 74.5 | CH | 3.81 | dd | 10.1 Hz (C4), 2.5 Hz (C6) | 5.33 ($^{3}J$, C1), 3.30 ($^{2}J$, C4), 1.13 ($^{3}J$, C7) |
| | 6 | 65.6 | CH | 4.12 | dd | 6.6 Hz (C7), 2.5 Hz (C5) | 3.81 ($^{2}J$, C2), 3.30 ($^{3}J$, C4), 1.13 ($^{2}J$, C7) |
| | 7 | 14.7 | $CH_3$ | 1.13 | d | 6.6 Hz (C6) | 14.7 ($^{1}J$, 127 Hz, C7), 3.81 ($^{3}J$, C5), 4.12 ($^{2}J$, C6) |

#### Experimental:

All NMR spectra were recorded at a Bruker Fourier spectrometer operating at 400.13 MHz ($^1H$) and 100.62 Hz $^{13}C$) respectively.
A sample of MCC3436 was dissolved in $D_2O$ and measured at 296 K.
$^1H$-NMR spectra were recorded using an excitation pulse of 30 degrees and a repetition time of 6.4 s.
16 scans were added and Fourier transformed with a final digital resolution of 0.18 Hz. The $^1H$-decoupled $^{13}C$ spectrum was recorded by adding 1024 transitions with an excitation pulse of 30 degrees and a repetition time of 3.35 s. Power gated decoupling was used to minimize sample heating. The spectral width was 24 kHz (320 ppm) with a digital resolution of 0.74 Hz.

The heteronuclear correlation spectrum (hsqc) was recorded by a matrix of 2k data points (f2, $^1H$ dimension) and 128 increments (data points f1 $^{13}C$ dimension).
The spectral width was 10 x 222 ppm.

A hetero-nuclear correlation via long range couplings (hmbc) was recorded with a data matrix of 2k data points (f2, $^1H$ dimension) and 512 increments (data points in f1 $^{13}C$ dimension). 8 scans were added for every time-increment resulting in an experiment time of 2 h. The spectral width was 15 x 160 ppm. with a relaxation delay of 1.5 s. A homo-nuclear correlation (cosy) was recorded with a data matrix of 2k data points (f2, $^1H$ dimension) and 256 increments (data points in f1). 4 scans were added for every time-increment resulting in an experiment time of 19 min. The spectral width was 13.5 ppm and the digital resolution in the indirect dimension was limited to 42 Hz.

A homo-nuclear correlation (cosy) was recorded with a data matrix of 2k data points (f2, $^1H$ dimension) and 256 increments (data points in f1). 2 scans were added for every time-increment resulting in an experiment time of 11 min. The spectral width was 13 ppm and the digital resolution in the indirect dimension was limited to 42 Hz.

Spectra were assigned using $^1H$-$^{13}C$ (hsqc and hmbc) 2D correlation spectra.

*Table 2: Calibration. Absolute referencing was used for 13C calibration.*

| CALIBRATION | $^1H$ |
|---|---|
| | HDO= 4.79 ppm |

Gentamicin B1 MCC

micro combichem
**MCC3436 RegID8096**
**Gentamicin B1**
**496.55g/mol**
Purity: Min. 97% Quantity: 5.6g
Long term Storage: -20°C
MicroCombiChem,65203 Wiesbaden Germany

$^1$H NMR (300 MHz, Deuterium Oxide) δ 5.61 (d, $J$ = 4.0 Hz, 1H), 5.12 (d, $J$ = 3.7 Hz, 1H), 4.26 – 4.15 (m, 2H), 4.04 – 3.76 (m, 7H), 3.62 – 3.38 (m, 7H), 3.30 (s, 1H), 2.89 (s, 3H), 2.53 (dt, $J$ = 12.4, 4.1 Hz, 1H), 1.94 (q, $J$ = 12.6 Hz, 1H), 1.31 (s, 3H), 1.18 (d, $J$ = 6.6 Hz, 3H).

$^{13}$C NMR (75 MHz, $D_2O$) δ 100.93, 97.28, 83.18, 81.37, 75.13, 73.50, 70.07, 69.84, 69.31, 67.67, 66.16, 65.17, 63.27, 54.06, 49.47, 48.84, 48.79, 34.46, 27.79, 20.81, 14.32.

**MCC3436 RegID8096**
**Gentamicin B1**
**496.55g/mol**
Purity: Min. 97% Quantity: 5.6g
Long term Storage: -20°C
MicroCombiChem,65203 Wiesbaden Germany

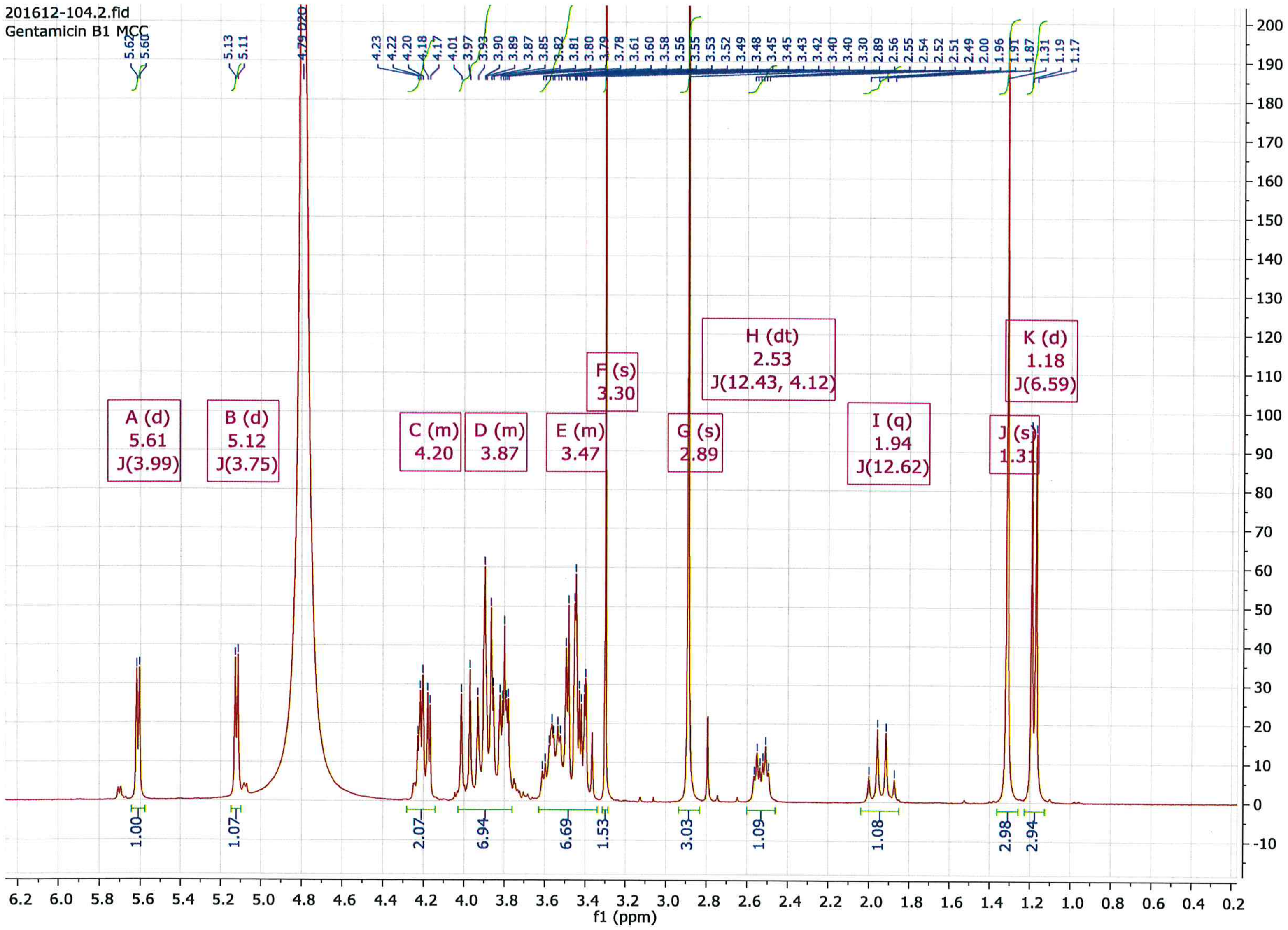

1:Proton

**MCC3436 RegID8096**
**Gentamicin B1**
**496.55g/mol**
Purity: Min. 97% Quantity: 5.6g
Long term Storage: -20°C
MicroCombiChem,65203 Wiesbaden Germany

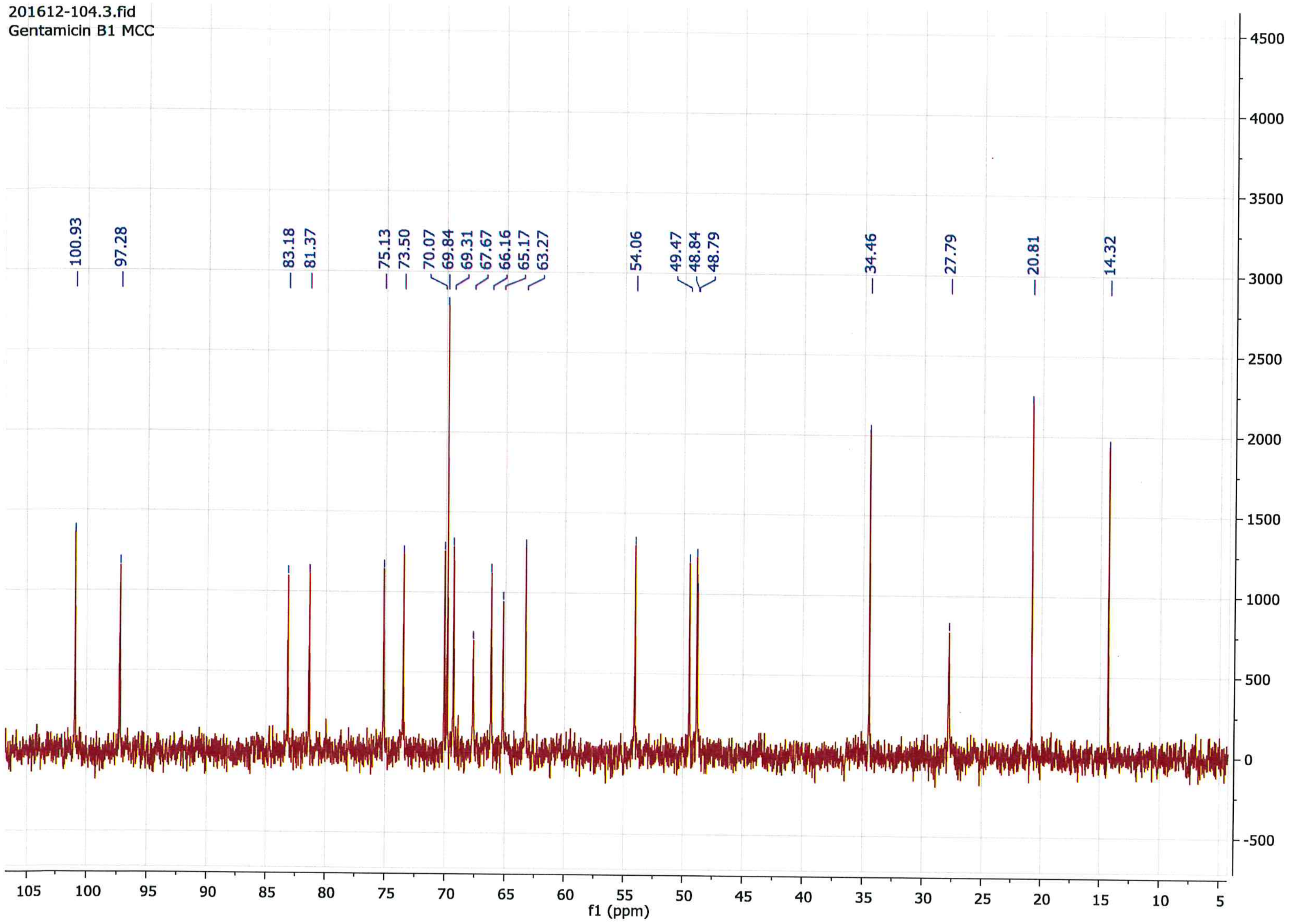

2:Carbon

**MCC3436 RegID8096**
**Gentamicin B1**
**496.55g/mol**
Purity: Min. 97% Quantity: 5.6g
Long term Storage: -20°C
MicroCombiChem,65203 Wiesbaden Germany

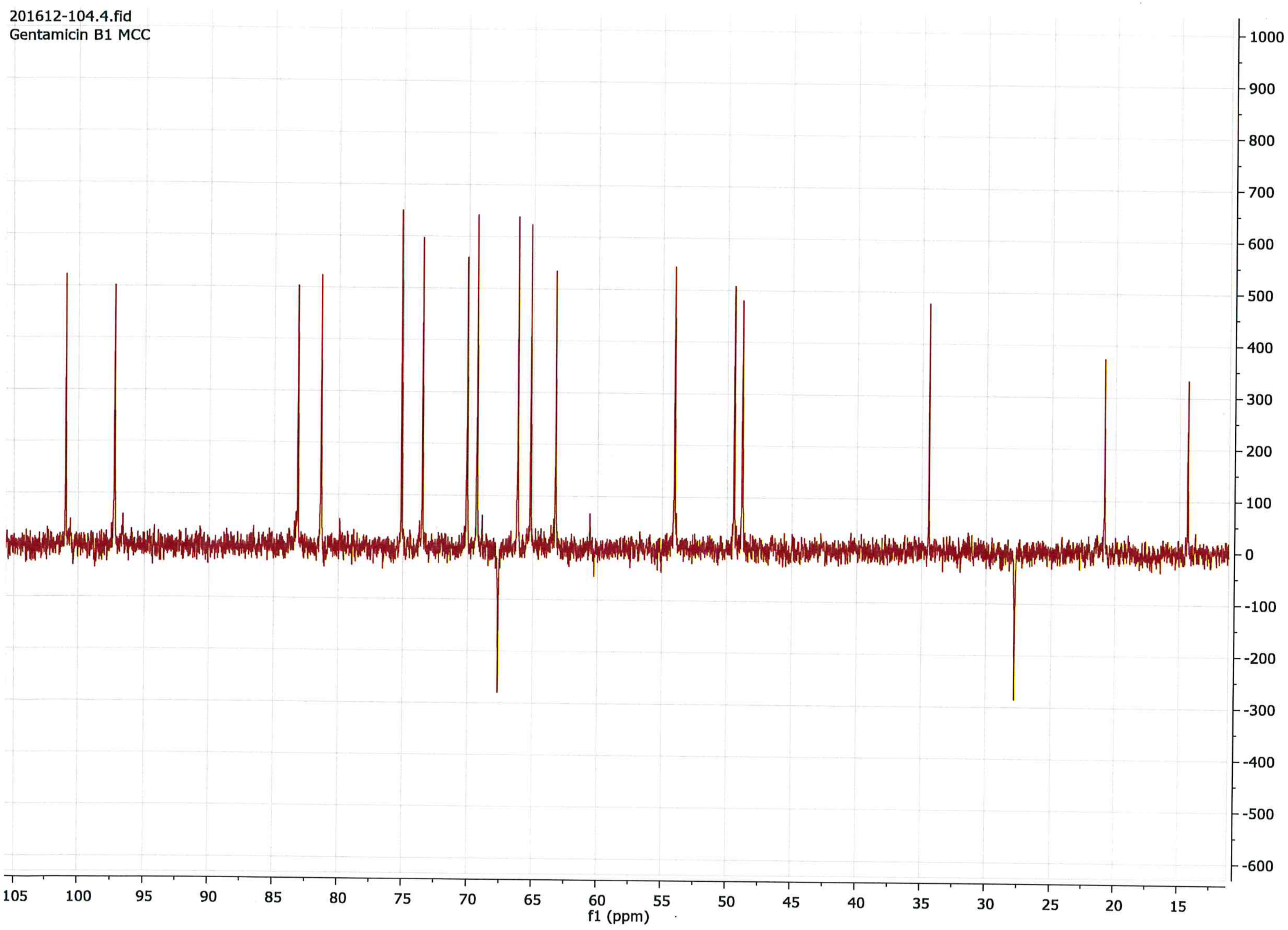

3:DEPT135

MCC3436 RegID8096
Gentamicin B1
496.55g/mol
Purity: Min. 97% Quantity: 5.6g
Long term Storage: -20°C
MicroCombiChem,65203 Wiesbaden Germany

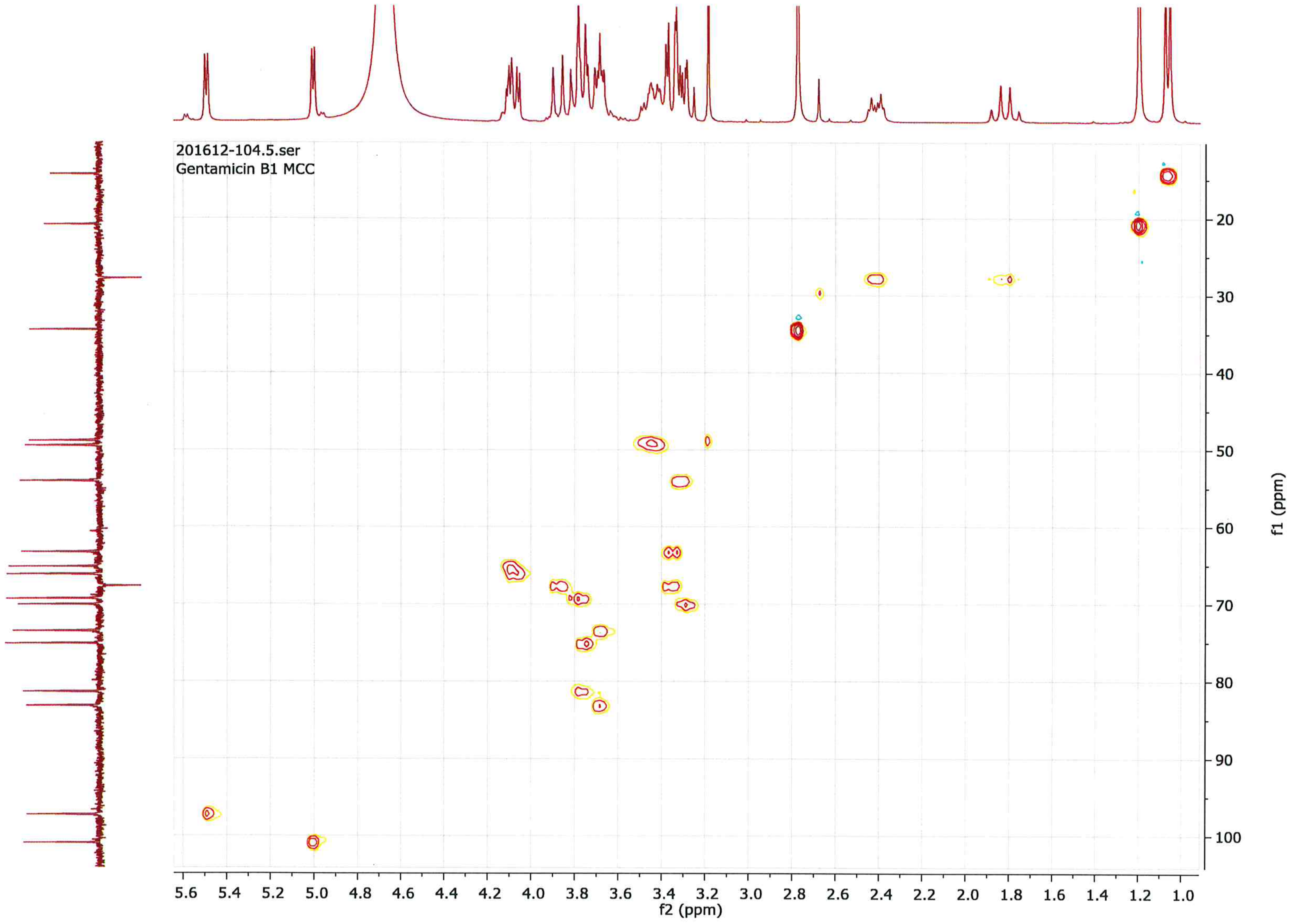

4:HSQC

**MCC3436 RegID8096**
**Gentamicin B1**
**496.55g/mol**
Purity: Min. 97% Quantity: 5.6g
Long term Storage: -20°C
MicroCombiChem,65203 Wiesbaden Germany

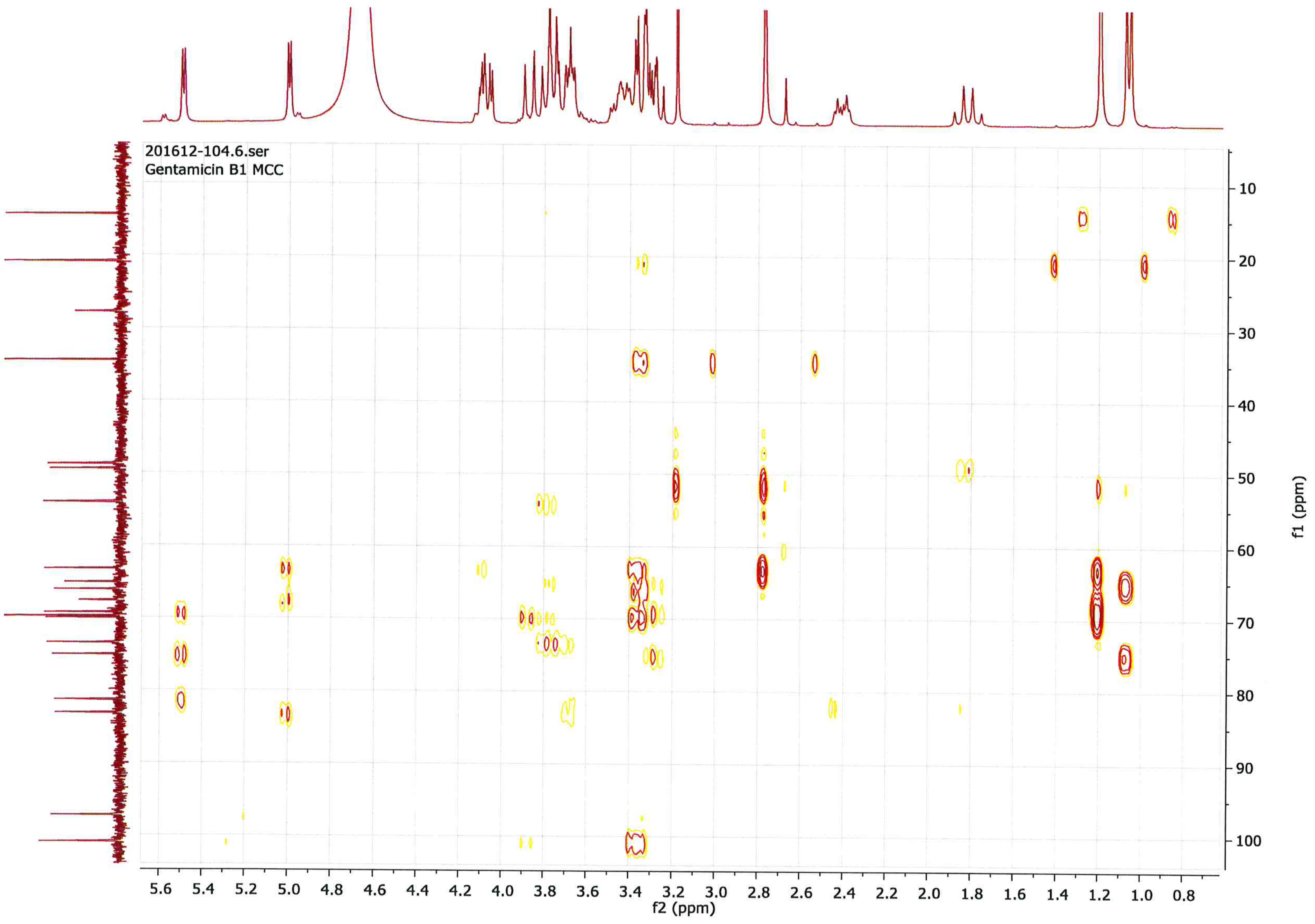

5:HMBC

**MCC3436 RegID8096**
**Gentamicin B1**
**496.55g/mol**
Purity: Min. 97% Quantity: 5.6g
Long term Storage: -20°C
MicroCombiChem,65203 Wiesbaden Germany

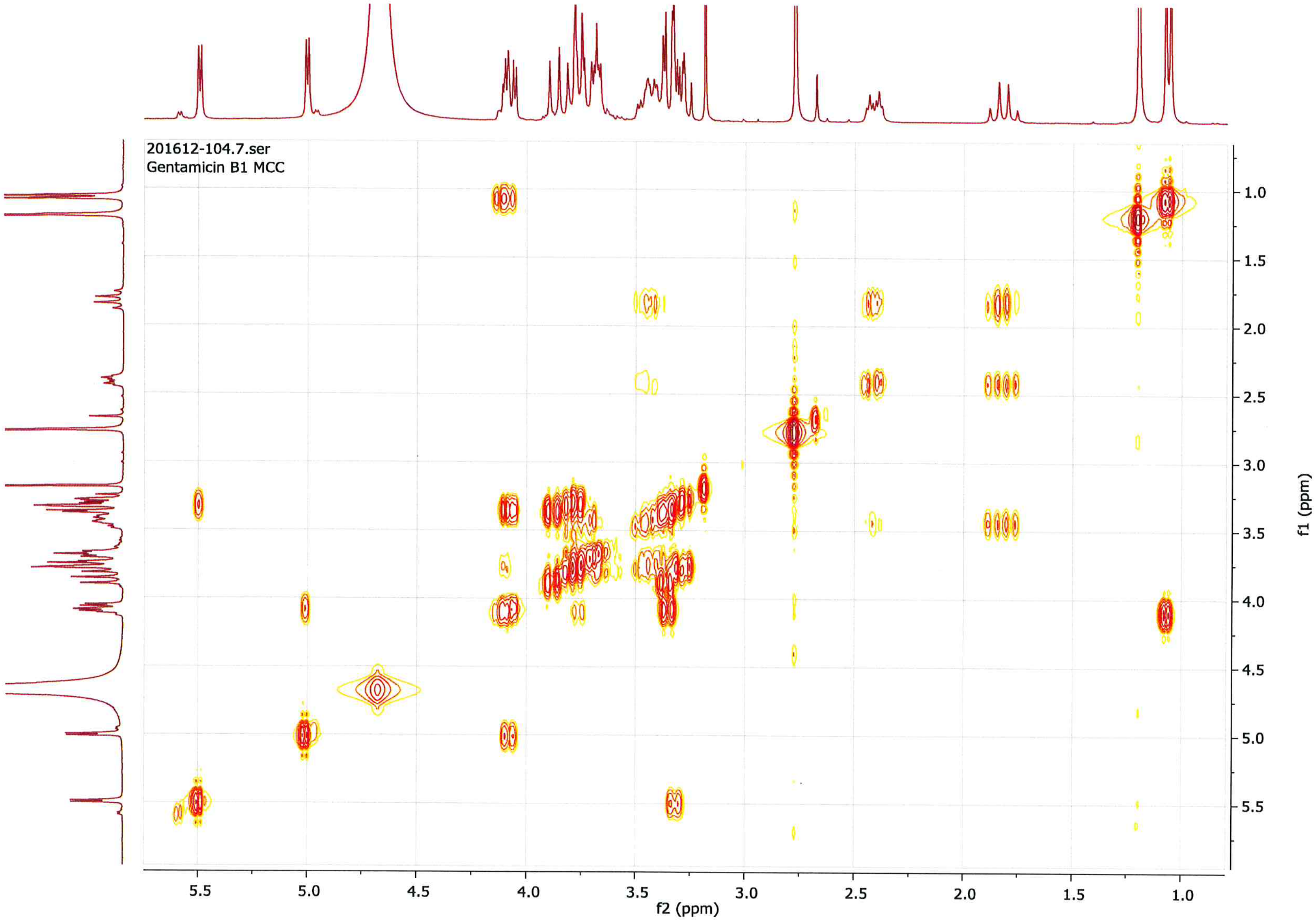

6: COSY
