## Supplementary material for "Gentamicin B1 is not a minor gentamicin component with major nonsense mutation suppression activity"

Toronto Research Chemicals
products for innovative research

### Certificate of Analysis

20 Martin Ross Ave., Toronto, ON. M3J 2K8 Canada Tel: (416) 665-9696 Fax: (416) 665-4439 Website: www.trc-canada.com

#### 1. Identification

**CAS Number:**
N/A

**Catalogue Number:**
G360560

**Product:**
Gentamicin B1 Acetate Salt

**Synonyms:**
Gentamycin B1 Acetate Salt; O-6-Amino-6,7-dideoxy-D-glycero-α-D-gluco-heptopyranosyl-(1→4)-O-[3-deoxy-4-C-methyl-3-(methylamino)-β-L-arabinopyranosyl-(1→6)]-2-deoxy-D-Streptamine Acetate Salt;

**Structure:**

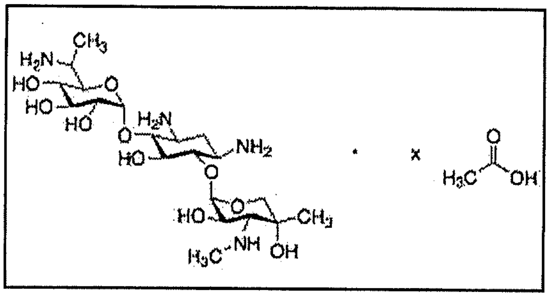

**Molecular Formula:**
$C_{20}H_{40}N_4O_{10} \cdot x(C_2H_4O_2)$

**Molecular Weight:**
496.55 + x(60.05)

**Source of Product:**
Synthetic

#### 2. Analytical Information

**Lot Number:**
9-MMS-23-1

**Melting Point:**
>166°C (dec.)

**Boiling Point:**
N/A

**Atmosphere:**
Inert Gas

**Appearance of Product:**
Off-White to Pale Yellow Solid

**Solubility**
Methanol (Very Slightly), Water (Sparingly)

**Method for Determining Identity:**
$^{1}H$ NMR ($D_2O$), $^{13}C$ NMR ($D_2O$), COSY ($D_2O$), HMBC ($D_2O$), HSQC ($D_2O$), and MS

**Stability**
Hygroscopic

**Purity:**
95%

**Long Term Storage Condition:**
Hygroscopic, Refrigerator, under inert atmosphere

**Additional Information:**
TLC Conditions: $C_{18}$; Water : Acetic Acid = 9 : 1; Visualized with AMCS and $KMnO_4$; Single Spot, Rf = 0.75.
$^{1}H$ NMR, $^{13}C$ NMR, COSY, HMBC, HSQC, and MS conform to structure.
Elemental Analysis: (Found) %C: 42.99, %H: 7.53, %N: 6.70; (Calculated) %C: 45.65, %H: 7.66, %N: 7.60 (for x = 4)
Acetate Content: 31.29% by Ion Chromatography

**Purity is based on the analytical results of the tests performed. NMR and TLC may have an accuracy of +/- 2%. Isotopic purity is based on mass distribution observed.**

*Philip Chan, Head of Quality Assurance*

**QC Test Date**
**July 4, 2018**

**Retest Date**
**July 2, 2022**

Toronto Research Chemicals
products for innovative research

**9-MMS-23-1 D2O**

```
Current Data Parameters
NAME      9-MMS-23-1, 20Jun2018
EXPNO                 10
PROCNO                 1

F2 - Acquisition Parameters
Date_           20180620
Time              22.43 h
INSTRUM            spect
PROBHD   Z116098_0683 (
PULPROG             zg30
TD                 32768
SOLVENT              D2O
NS                     8
DS                     1
SWH             7211.539 Hz
FIDRES          0.440157 Hz
AQ             2.2719147 sec
RG                  49.8
DW                69.333 usec
DE                  6.50 usec
TE                 298.0 K
D1            1.00000000 sec
TD0                    1
SFO1       400.3328023 MHz
NUC1                  1H
P1                 19.00 usec
PLW1          4.09749985 W

F2 - Processing parameters
SI                 32768
SF         400.3300000 MHz
WDW                   EM
SSB      0
LB                  0.30 Hz
GB       0
PC                 10.00
```

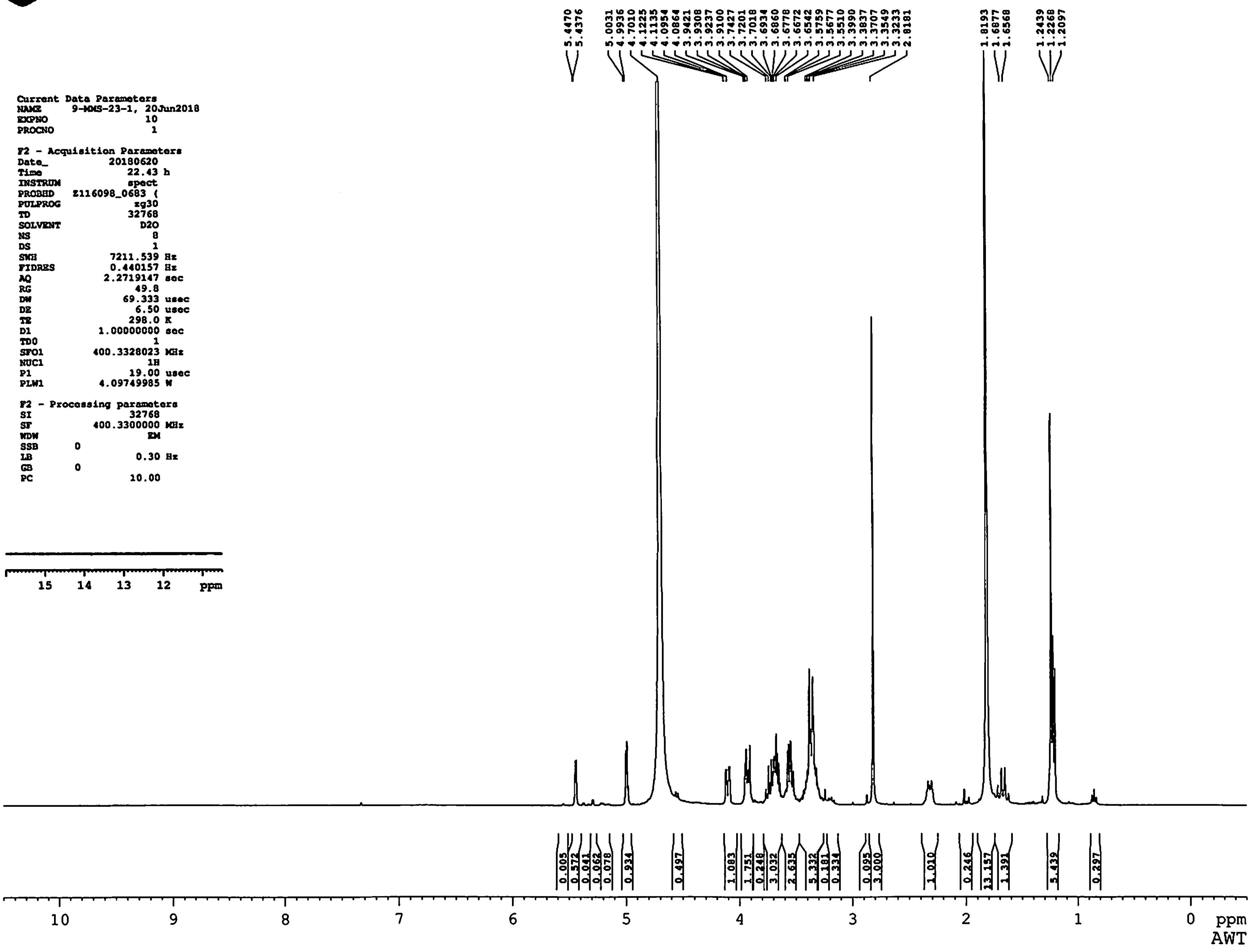

Toronto Research Chemicals
products for innovative research

**9-MMS-23-1 D2O**

```
Current Data Parameters
NAME      9-MMS-23-1, 8Jun2018
EXPNO                10
PROCNO                1

F2 - Acquisition Parameters
Date_          20180609
Time              14.57 h
INSTRUM           spect
PROBHD   Z116098_0683 (
PULPROG            zg30
TD                32768
SOLVENT             D2O
NS                  256
DS                    1
SWH            7211.539 Hz
FIDRES         0.440157 Hz
AQ            2.2719147 sec
RG               113.73
DW               69.333 usec
DE                 6.50 usec
TE                298.0 K
D1           1.00000000 sec
TD0                   1
SFO1        400.3328023 MHz
NUC1                 1H
P1                19.00 usec
PLW1         4.09749985 W

F2 - Processing parameters
SI                32768
SF          400.3300000 MHz
WDW                  EM
SSB      0
LB                 0.30 Hz
GB       0
PC                 5.00
```

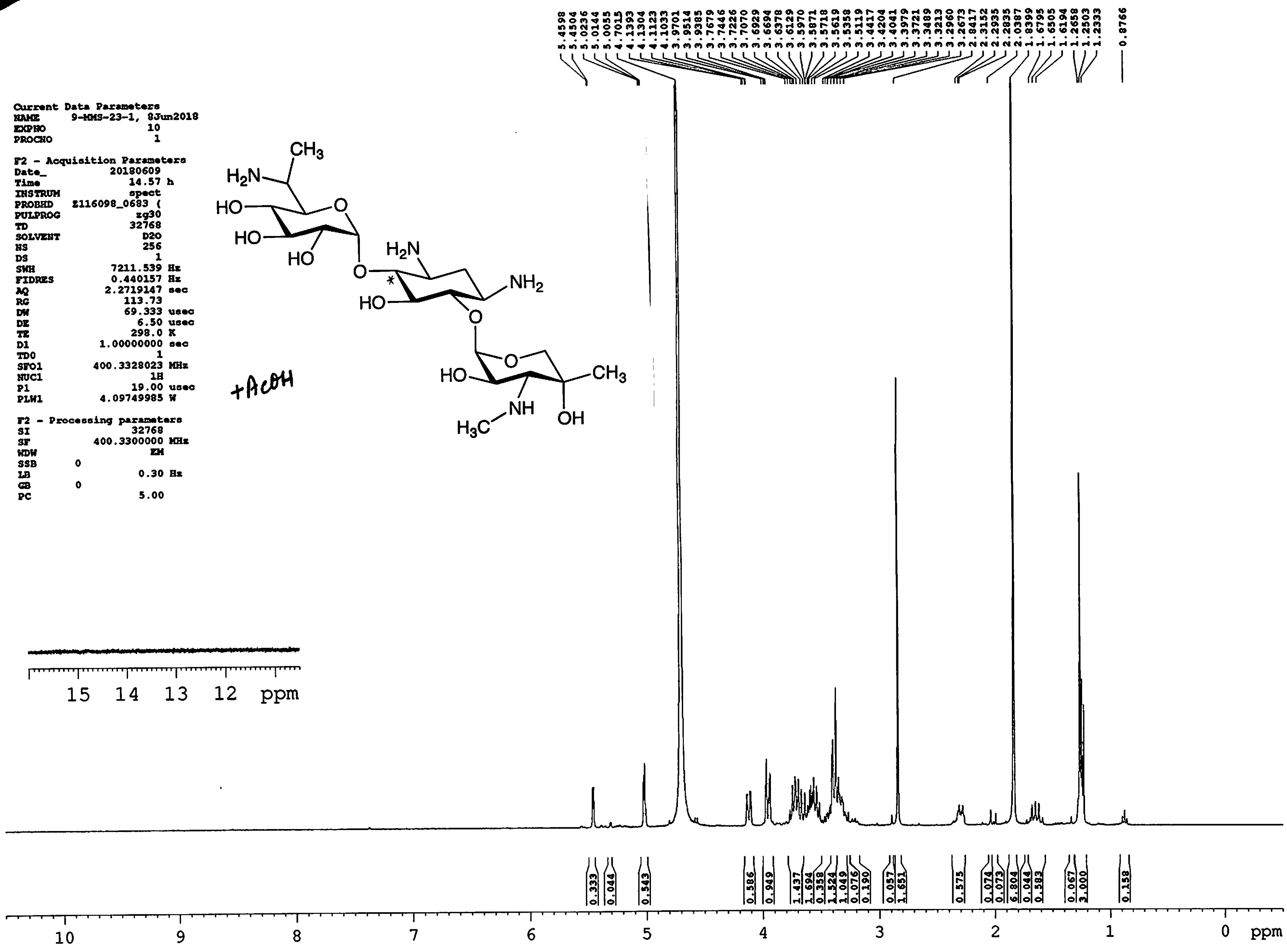

Toronto Research Chemicals
products for innovative research

### 9-MMS-23-1 D2O 13C

104.374
101.032
100.881
99.001
95.580
84.619
83.963
83.665
79.485
74.625
73.767
72.725
72.401
72.358
70.858
70.787
69.897
69.863
69.703
67.527
66.311
63.529
50.119
49.992
49.827
47.519
47.051
34.534
29.042
28.359
23.171
20.849
11.110
10.944
8.477

| | |
|---|---|
| **Current Data Parameters** | |
| NAME | 9-MMS-23-1, 20Jun2018 |
| EXPNO | 11 |
| PROCNO | 1 |
| **F2 - Acquisition Parameters** | |
| Date_ | 20180621 |
| Time | 1.40 h |
| INSTRUM | spect |
| PROBHD | Z116098_0683 ( |
| PULPROG | zgpg30 |
| TD | 65536 |
| SOLVENT | D2O |
| NS | 3072 |
| DS | 4 |
| SWH | 24038.461 Hz |
| FIDRES | 0.733596 Hz |
| AQ | 1.3631488 sec |
| RG | 203.4 |
| DW | 20.800 usec |
| DE | 6.50 usec |
| TE | 298.0 K |
| D1 | 2.00000000 sec |
| D11 | 0.03000000 sec |
| TD0 | 1 |
| SFO1 | 100.6731249 MHz |
| NUC1 | 13C |
| P1 | 10.00 usec |
| PLW1 | 66.69200134 W |
| SFO2 | 400.3316013 MHz |
| NUC2 | 1H |
| CPDPRG[2 | waltz16 |
| PCPD2 | 90.00 usec |
| PLW2 | 4.09749985 W |
| PLW12 | 0.18262000 W |
| PLW13 | 0.09185500 W |
| **F2 - Processing parameters** | |
| SI | 32768 |
| SF | 100.6630586 MHz |
| WDW | EM |
| SSB | 0 |
| LB | 1.00 Hz |
| GB | 0 |
| PC | 1.40 |

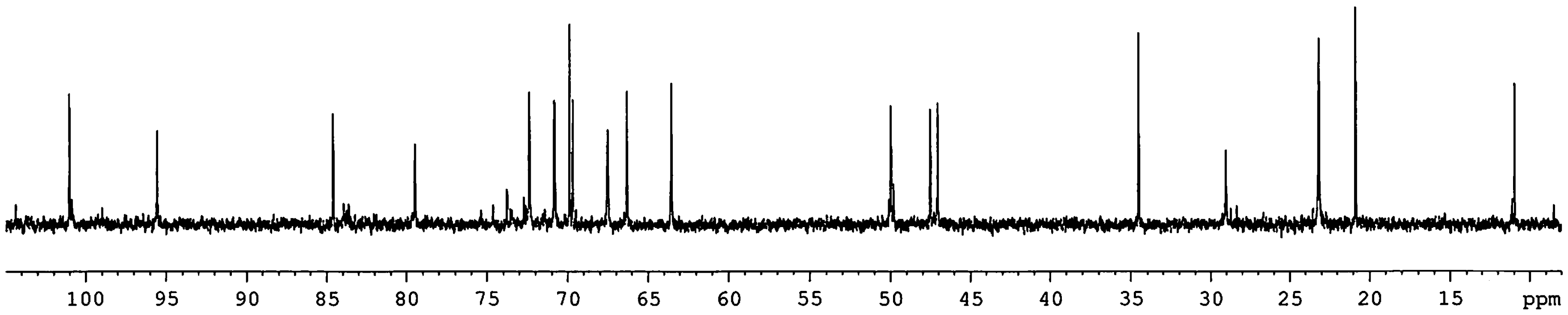


AWT

*9-MMS-23-1 D2O edHSQC*

Toronto Research Chemicals
products for innovative research

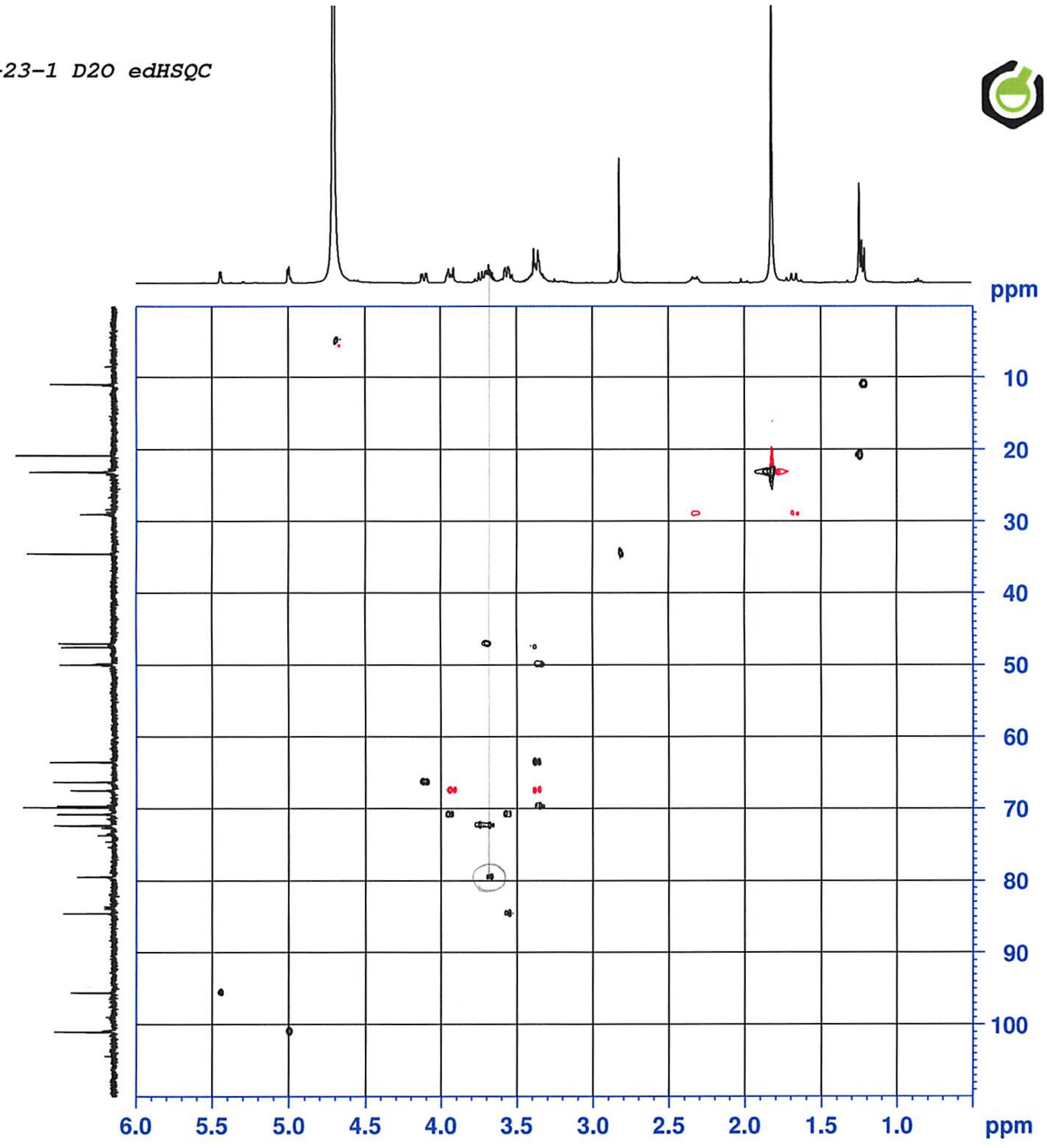


```
Current Data Parameters
NAME       9-MMS-23-1, 20Jun2018
EXPNO                      12
PROCNO                      1

F2 - Acquisition Parameters
Date_                20180621
Time                     1.42 h
INSTRUM                 spect
PROBHD     Z116098_0683 (
PULPROG    hsqcedetgpsp.3
TD                       2048
SOLVENT                   D2O
NS                          4
DS                         16
SWH                  4807.692 Hz
FIDRES               4.695012 Hz
AQ                  0.2129920 sec
RG                      203.4
DW                    104.000 usec
DE                       6.50 usec
TE                      298.0 K
CNST2             145.0000000
D0                 0.00000300 sec
D1                 2.00000000 sec
D4                 0.00172414 sec
D11                0.03000000 sec
D16                0.00020000 sec
D21                0.00360000 sec
IN0                0.00002760 sec
TDav                        1
SFO1              400.3324020 MHz
NUC1                       1H
P1                      19.00 usec
P2                      38.00 usec
P28                   1000.00 usec
PLW1               4.09749985 W
SFO2              100.6721182 MHz
NUC2                      13C
CPDPRG[2 bi_p5m4sp_4sp.2
P3                      10.00 usec
P14                    500.00 usec
P31                   2119.00 usec
P63                   1500.00 usec
PLW0       0 W
PLW2              66.69200134 W
PLW12              1.04209995 W
SPNAM[3] Crp60,0.5,20.1
SPOAL3                  0.500
SPOFFS3    0 Hz
SPW3              10.18999958 W
SPNAM[14 Crp32,1.5,20.2
SPOAL14                 0.500
SPOFFS14 0 Hz
SPW14              4.34770012 W
SPNAM[18  Crp60_xfilt.2
SPOAL18                 0.500
SPOFFS18 0 Hz
SPW18              1.96300006 W
SPNAM[31 Crp32,1.5,20.2
SPOAL31                 0.500
SPOFFS31 0 Hz
SPW31              1.08690000 W
GPNAM[1]           SMSQ10.100
GPZ1                    80.00 %
GPNAM[2]           SMSQ10.100
GPZ2                    20.10 %
P16                   1000.00 usec

F1 - Acquisition parameters
TD                        256
SFO1                 100.6721 MHz
FIDRES             141.530792 Hz
SW                    179.950 ppm
FnMODE         Echo-Antiecho

F2 - Processing parameters
SI                       2048
SF                400.3300041 MHz
WDW                     QSINE
SSB                         2
LB         0 Hz
GB         0
PC                       1.40

F1 - Processing parameters
SI                       1024
MC2            echo-antiecho
SF                100.6630586 MHz
WDW                     QSINE
SSB                         2
LB         0 Hz
GB         0
```

*9-MMS-23-1 D2O COSY*

Toronto Research Chemicals
products for innovative research

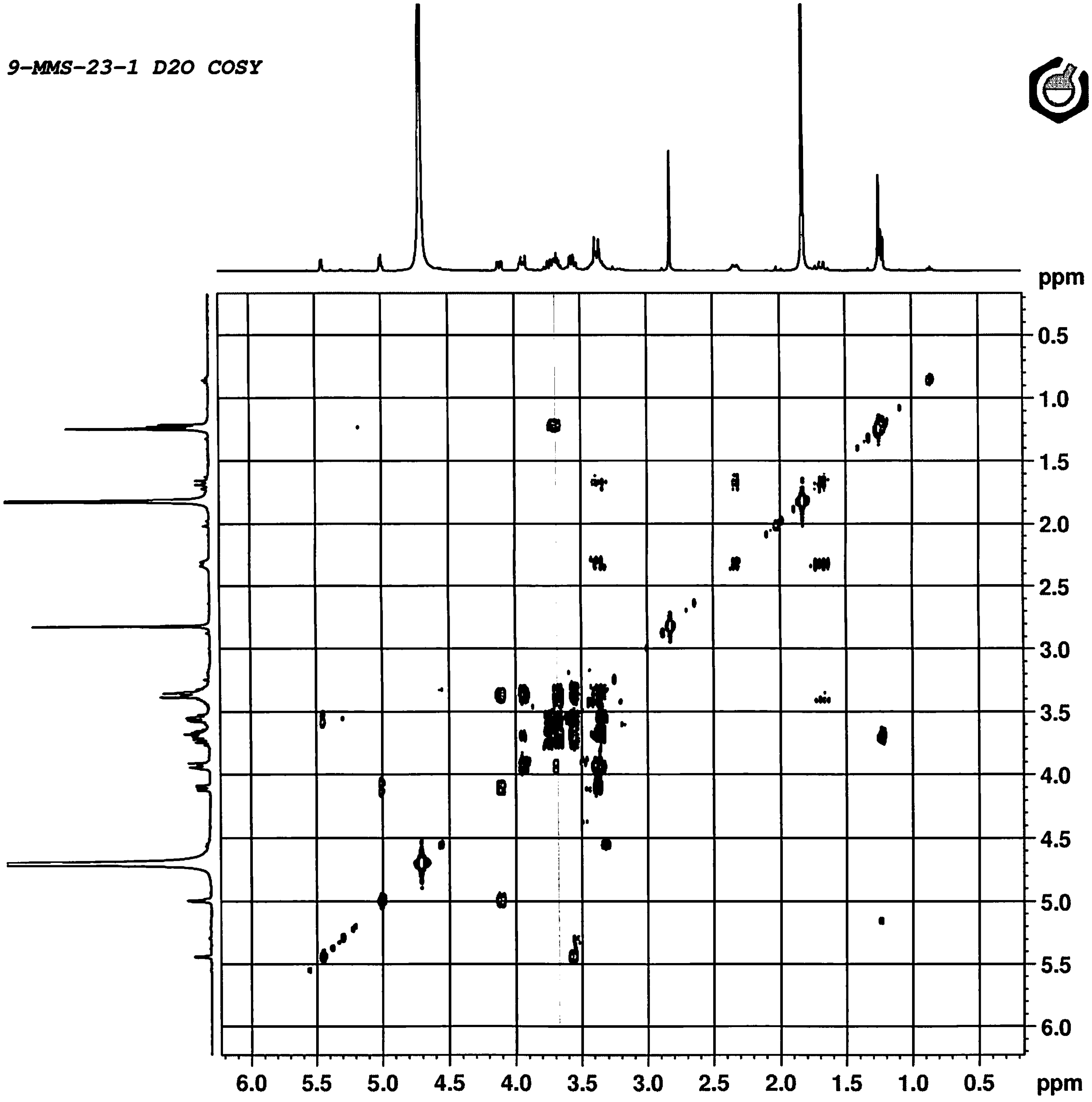


**Current Data Parameters**

| | |
|---|---|
| NAME | 9-MMS-23-1, 20Jun2018 |
| EXPNO | 15 |
| PROCNO | 1 |

**F2 - Acquisition Parameters**

| | |
|---|---|
| Date_ | 20180621 |
| Time | 4.35 h |
| INSTRUM | spect |
| PROBHD | Z116098_0683 ( |
| PULPROG | cosygpppqf |
| TD | 2048 |
| SOLVENT | D2O |
| NS | 1 |
| DS | 16 |
| SWH | 2427.185 Hz |
| FIDRES | 2.370297 Hz |
| AQ | 0.4218880 sec |
| RG | 35.31 |
| DW | 206.000 usec |
| DE | 6.50 usec |
| TE | 298.0 K |
| D0 | 0.00000300 sec |
| D1 | 1.77471995 sec |
| D11 | 0.03000000 sec |
| D12 | 0.00002000 sec |
| D13 | 0.00000400 sec |
| D16 | 0.00020000 sec |
| IN0 | 0.00041200 sec |
| TDav | 1 |
| SFO1 | 400.3312801 MHz |
| NUC1 | 1H |
| P0 | 19.00 usec |
| P1 | 19.00 usec |
| P17 | 2500.00 usec |
| PLW1 | 4.09749985 W |
| PLW10 | 1.64359999 W |
| GPNAM[1] | SMSQ10.100 |
| GPZ1 | 10.00 % |
| P16 | 1000.00 usec |

**F1 - Acquisition parameters**

| | |
|---|---|
| TD | 128 |
| SFO1 | 400.3313 MHz |
| FIDRES | 37.924759 Hz |
| SW | 6.063 ppm |
| FnMODE | QF |

**F2 - Processing parameters**

| | |
|---|---|
| SI | 1024 |
| SF | 400.3300000 MHz |
| WDW | QSINE |
| SSB | 0 |
| LB | 0 Hz |
| GB | 0 |
| PC | 1.40 |

**F1 - Processing parameters**

| | |
|---|---|
| SI | 1024 |
| MC2 | QF |
| SF | 400.3300000 MHz |
| WDW | QSINE |
| SSB | 0 |
| LB | 0 Hz |
| GB | 0 |

Toronto Research Chemicals
products for innovative research

**9-MMS-23-1 D2O 13C**

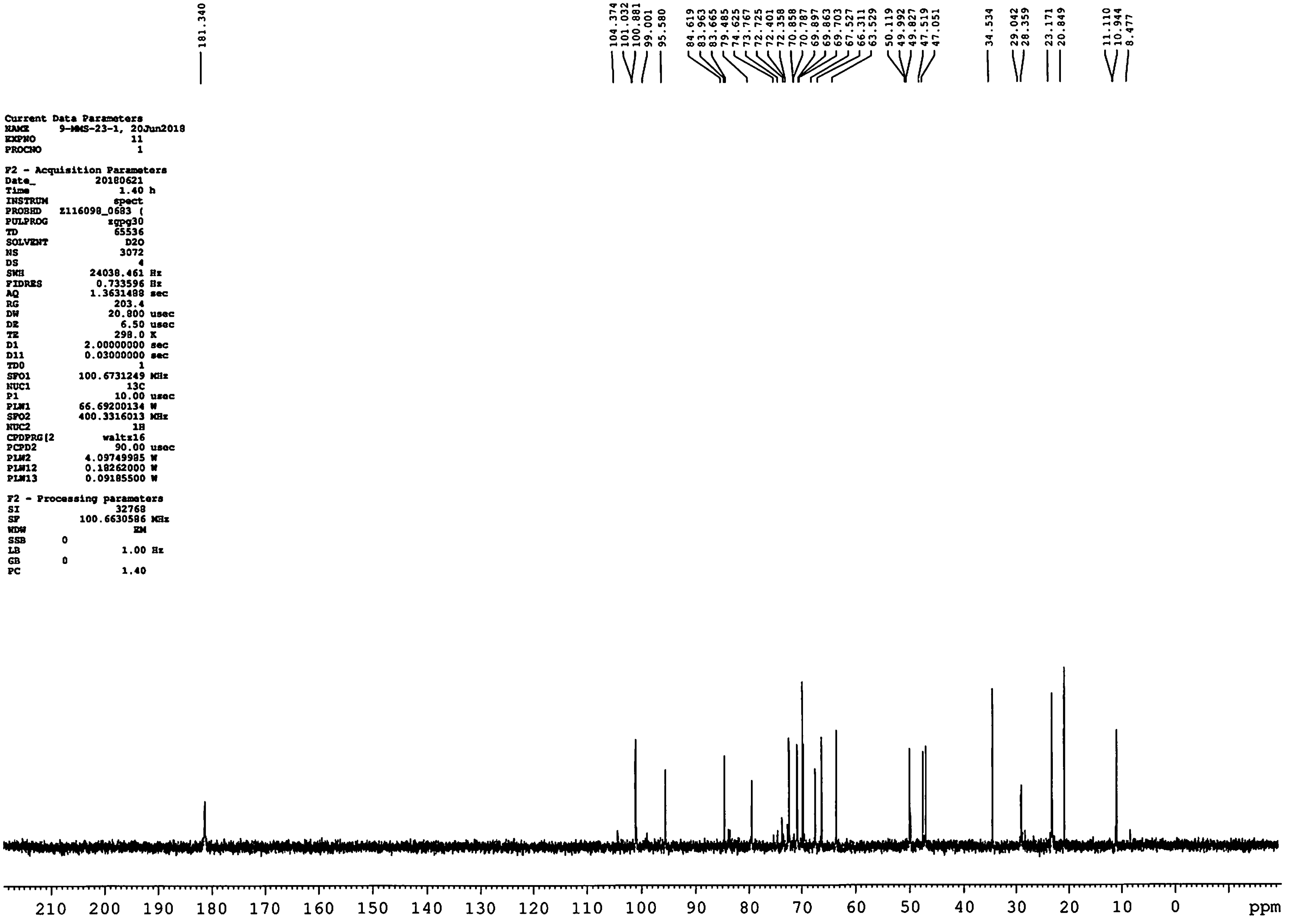


Current Data Parameters
NAME 9-MMS-23-1, 20Jun2018
EXPNO 11
PROCNO 1

F2 - Acquisition Parameters
Date_ 20180621
Time 1.40 h
INSTRUM spect
PROBHD Z116098_0683 (
PULPROG zgpg30
TD 65536
SOLVENT D2O
NS 3072
DS 4
SWH 24038.461 Hz
FIDRES 0.733596 Hz
AQ 1.3631488 sec
RG 203.4
DW 20.800 usec
DE 6.50 usec
TE 298.0 K
D1 2.00000000 sec
D11 0.03000000 sec
TD0 1
SFO1 100.6731249 MHz
NUC1 13C
P1 10.00 usec
PLW1 66.69200134 W
SFO2 400.3316013 MHz
NUC2 1H
CPDPRG[2 waltz16
PCPD2 90.00 usec
PLW2 4.09749985 W
PLW12 0.18262000 W
PLW13 0.09185500 W

F2 - Processing parameters
SI 32768
SF 100.6630586 MHz
WDW EM
SSB 0
LB 1.00 Hz
GB 0
PC 1.40

AWT

*9-MMS-23-1 D2O edHSQC*

Toronto Research Chemicals
products for innovative research

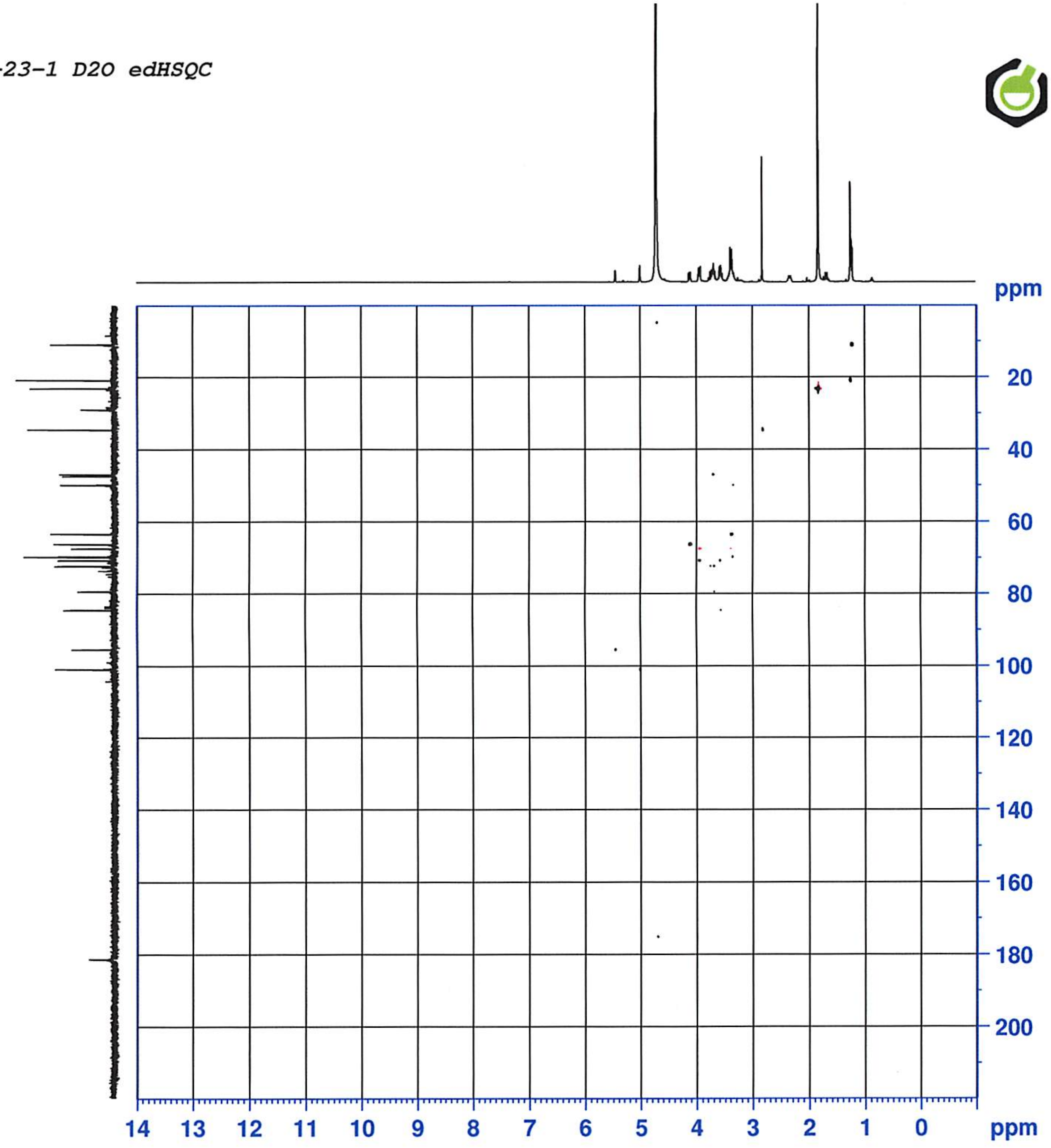


```
Current Data Parameters
NAME      9-MMS-23-1, 20Jun2018
EXPNO                    12
PROCNO                    1

F2 - Acquisition Parameters
Date_              20180621
Time                 1.42 h
INSTRUM               spect
PROBHD    Z116098_0683 (
PULPROG   hsqcedetgpsp.3
TD                     2048
SOLVENT                 D2O
NS                        4
DS                       16
SWH                4807.692 Hz
FIDRES             4.695012 Hz
AQ                0.2129920 sec
RG                    203.4
DW                  104.000 usec
DE                     6.50 usec
TE                    298.0 K
CNST2           145.0000000
D0               0.00000300 sec
D1               2.00000000 sec
D4               0.00172414 sec
D11              0.03000000 sec
D16              0.00020000 sec
D21              0.00360000 sec
IN0              0.00002760 sec
TDav                      1
SFO1            400.3324020 MHz
NUC1                     1H
P1                    19.00 usec
P2                    38.00 usec
P28                 1000.00 usec
PLW1             4.09749985 W
SFO2            100.6721182 MHz
NUC2                    13C
CPDPRG[2 bi_p5m4sp_4sp.2
P3                    10.00 usec
P14                  500.00 usec
P31                 2119.00 usec
P63                 1500.00 usec
PLW0      0 W
PLW2            66.69200134 W
PLW12            1.04209995 W
SPNAM[3] Crp60,0.5,20.1
SPOAL3                0.500
SPOFFS3   0 Hz
SPW3            10.18999958 W
SPNAM[14 Crp32,1.5,20.2
SPOAL14               0.500
SPOFFS14 0 Hz
SPW14            4.34770012 W
SPNAM[18  Crp60_xfilt.2
SPOAL18               0.500
SPOFFS18 0 Hz
SPW18            1.96300006 W
SPNAM[31 Crp32,1.5,20.2
SPOAL31               0.500
SPOFFS31 0 Hz
SPW31            1.08690000 W
GPNAM[1]      SMSQ10.100
GPZ1                  80.00 %
GPNAM[2]      SMSQ10.100
GPZ2                  20.10 %
P16                 1000.00 usec

F1 - Acquisition parameters
TD                      256
SFO1               100.6721 MHz
FIDRES           141.530792 Hz
SW                  179.950 ppm
FnMODE     Echo-Antiecho

F2 - Processing parameters
SI                     2048
SF              400.3300041 MHz
WDW                   QSINE
SSB                       2
LB        0 Hz
GB        0
PC                     1.40

F1 - Processing parameters
SI                     1024
MC2          echo-antiecho
SF              100.6630586 MHz
WDW                   QSINE
SSB                       2
LB        0 Hz
GB        0
```

*9-MMS-23-1 D2O HMBC*

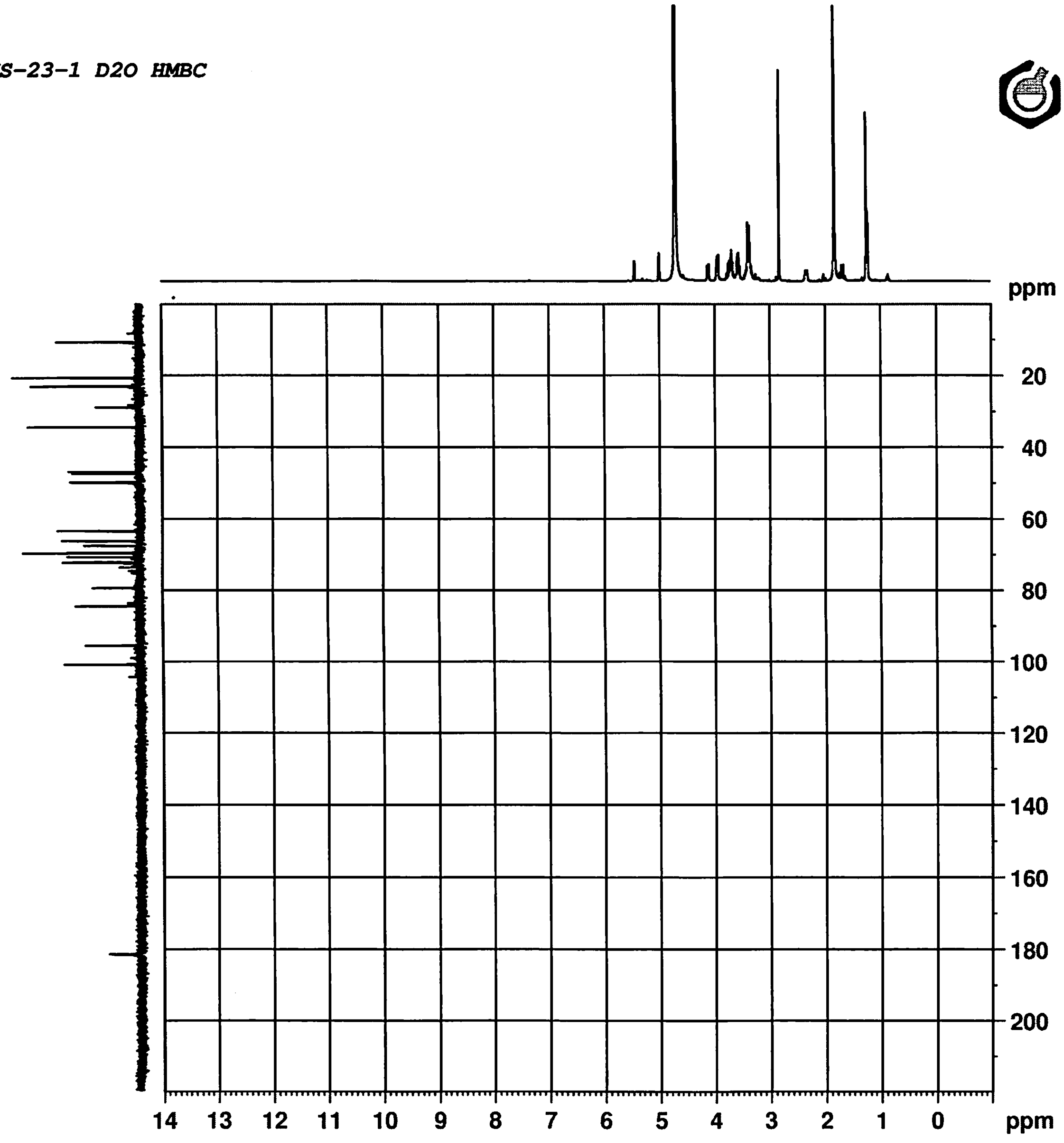


Toronto Research Chemicals
products for innovative research

**Current Data Parameters**

| | |
|---|---|
| NAME | 9-MMS-23-1, 20Jun2018 |
| EXPNO | 13 |
| PROCNO | 1 |

**F2 - Acquisition Parameters**

| | |
|---|---|
| Date_ | 20180621 |
| Time | 2.22 h |
| INSTRUM | spect |
| PROBHD | Z116098_0683 ( |
| PULPROG | hmbcetgpl3nd |
| TD | 4096 |
| SOLVENT | D2O |
| NS | 8 |
| DS | 16 |
| SWH | 6009.615 Hz |
| FIDRES | 2.934382 Hz |
| AQ | 0.3407872 sec |
| RG | 203.4 |
| DW | 83.200 usec |
| DE | 6.50 usec |
| TE | 297.9 K |
| CNST6 | 120.0000000 |
| CNST7 | 170.0000000 |
| CNST13 | 8.0000000 |
| D0 | 0.00000300 sec |
| D1 | 1.50000000 sec |
| D6 | 0.06250000 sec |
| D16 | 0.00020000 sec |
| IN0 | 0.00002260 sec |
| TDav | 1 |
| SFO1 | 400.3326021 MHz |
| NUC1 | 1H |
| P1 | 19.00 usec |
| P2 | 38.00 usec |
| PLW1 | 4.09749985 W |
| SFO2 | 100.6741315 MHz |
| NUC2 | 13C |
| P3 | 10.00 usec |
| P24 | 2000.00 usec |
| PLW2 | 66.69200134 W |
| SPNAM[7] | Crp60comp.4 |
| SPOAL7 | 0.500 |
| SPOFFS7 | 0 Hz |
| SPW7 | 10.18999958 W |
| GPNAM[1] | SMSQ10.100 |
| GPZ1 | 80.00 % |
| GPNAM[3] | SMSQ10.100 |
| GPZ3 | 14.00 % |
| GPNAM[4] | SMSQ10.100 |
| GPZ4 | -8.00 % |
| GPNAM[5] | SMSQ10.100 |
| GPZ5 | -4.00 % |
| GPNAM[6] | SMSQ10.100 |
| GPZ6 | -2.00 % |
| P16 | 1000.00 usec |

**F1 - Acquisition parameters**

| | |
|---|---|
| TD | 512 |
| SFO1 | 100.6741 MHz |
| FIDRES | 86.421463 Hz |
| SW | 219.757 ppm |
| FnMODE | Echo-Antiecho |

**F2 - Processing parameters**

| | |
|---|---|
| SI | 2048 |
| SF | 400.3300000 MHz |
| WDW | SINE |
| SSB | 0 |
| LB | 0 Hz |
| GB | 0 |
| PC | 1.40 |

**F1 - Processing parameters**

| | |
|---|---|
| SI | 2048 |
| MC2 | echo-antiecho |
| SF | 100.6630586 MHz |
| WDW | QSINE |
| SSB | 2 |
| LB | 0 Hz |
| GB | 0 |

9-MMS-23-1 D2O HMBC

Toronto Research Chemicals
products for innovative research

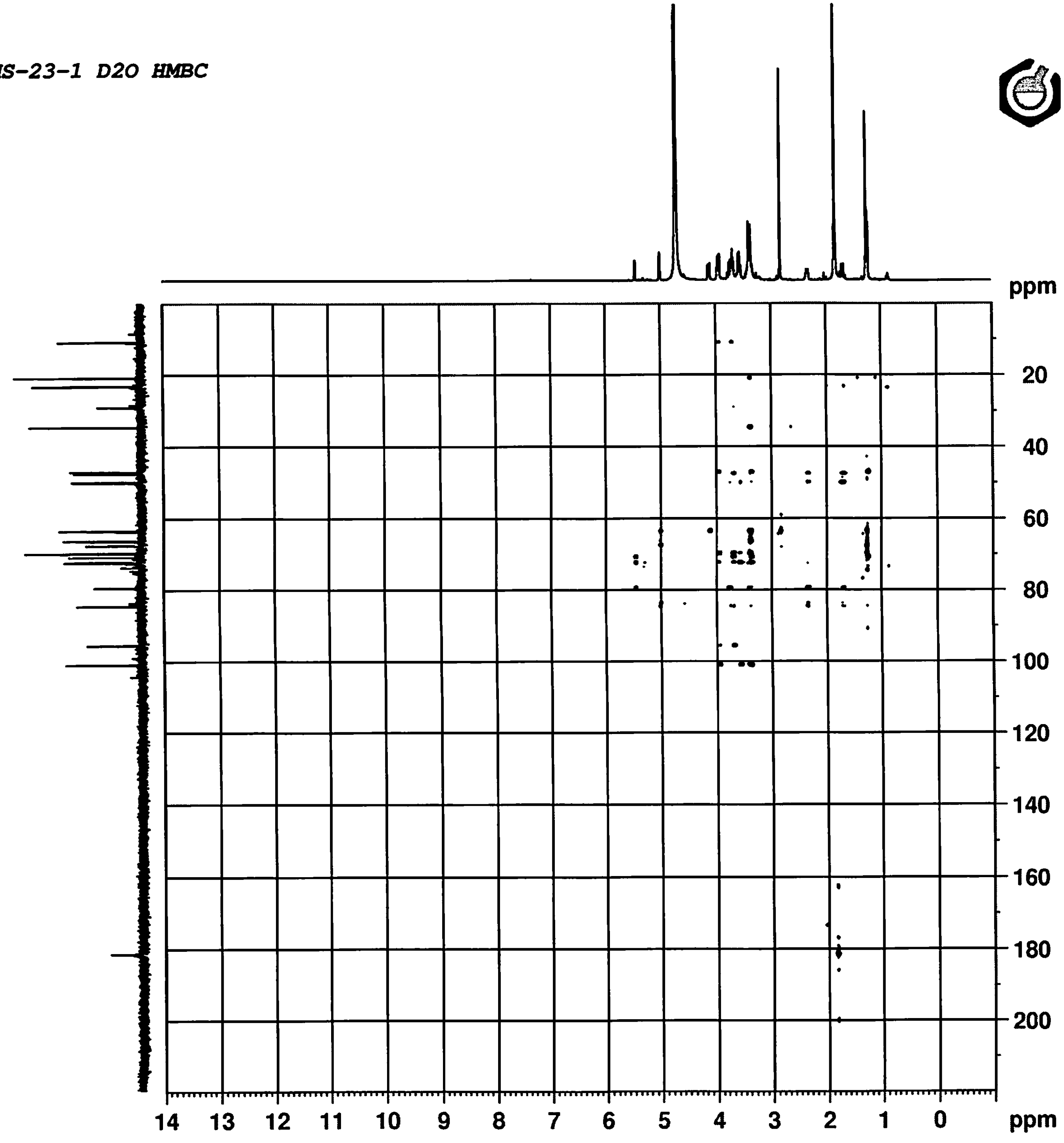


```
Current Data Parameters
NAME       9-MMS-23-1, 20Jun2018
EXPNO              13
PROCNO              1

F2 - Acquisition Parameters
Date_           20180621
Time                2.22 h
INSTRUM            spect
PROBHD    Z116098_0683 (
PULPROG      hmbcetgpl3nd
TD                  4096
SOLVENT              D2O
NS                     8
DS                    16
SWH             6009.615 Hz
FIDRES          2.934382 Hz
AQ             0.3407872 sec
RG                 203.4
DW                83.200 usec
DE                  6.50 usec
TE                 297.9 K
CNST6        120.0000000
CNST7        170.0000000
CNST13         8.0000000
D0            0.00000300 sec
D1            1.50000000 sec
D6            0.06250000 sec
D16           0.00020000 sec
IN0           0.00002260 sec
TDav                   1
SFO1         400.3326021 MHz
NUC1                  1H
P1                 19.00 usec
P2                 38.00 usec
PLW1          4.09749985 W
SFO2         100.6741315 MHz
NUC2                 13C
P3                 10.00 usec
P24              2000.00 usec
PLW2         66.69200134 W
SPNAM[7]    Crp60comp.4
SPOAL7             0.500
SPOFFS7  0 Hz
SPW7         10.18999958 W
GPNAM[1]      SMSQ10.100
GPZ1               80.00 %
GPNAM[3]      SMSQ10.100
GPZ3               14.00 %
GPNAM[4]      SMSQ10.100
GPZ4               -8.00 %
GPNAM[5]      SMSQ10.100
GPZ5               -4.00 %
GPNAM[6]      SMSQ10.100
GPZ6               -2.00 %
P16              1000.00 usec

F1 - Acquisition parameters
TD                   512
SFO1            100.6741 MHz
FIDRES         86.421463 Hz
SW               219.757 ppm
FnMODE      Echo-Antiecho

F2 - Processing parameters
SI                  2048
SF           400.3300000 MHz
WDW                 SINE
SSB         0
LB          0 Hz
GB          0
PC                  1.40

F1 - Processing parameters
SI                  2048
MC2          echo-antiecho
SF           100.6630586 MHz
WDW                QSINE
SSB                    2
LB          0 Hz
GB          0
```

*9-MMS-23-1 D2O HMBC*

Toronto Research Chemicals
products for innovative research

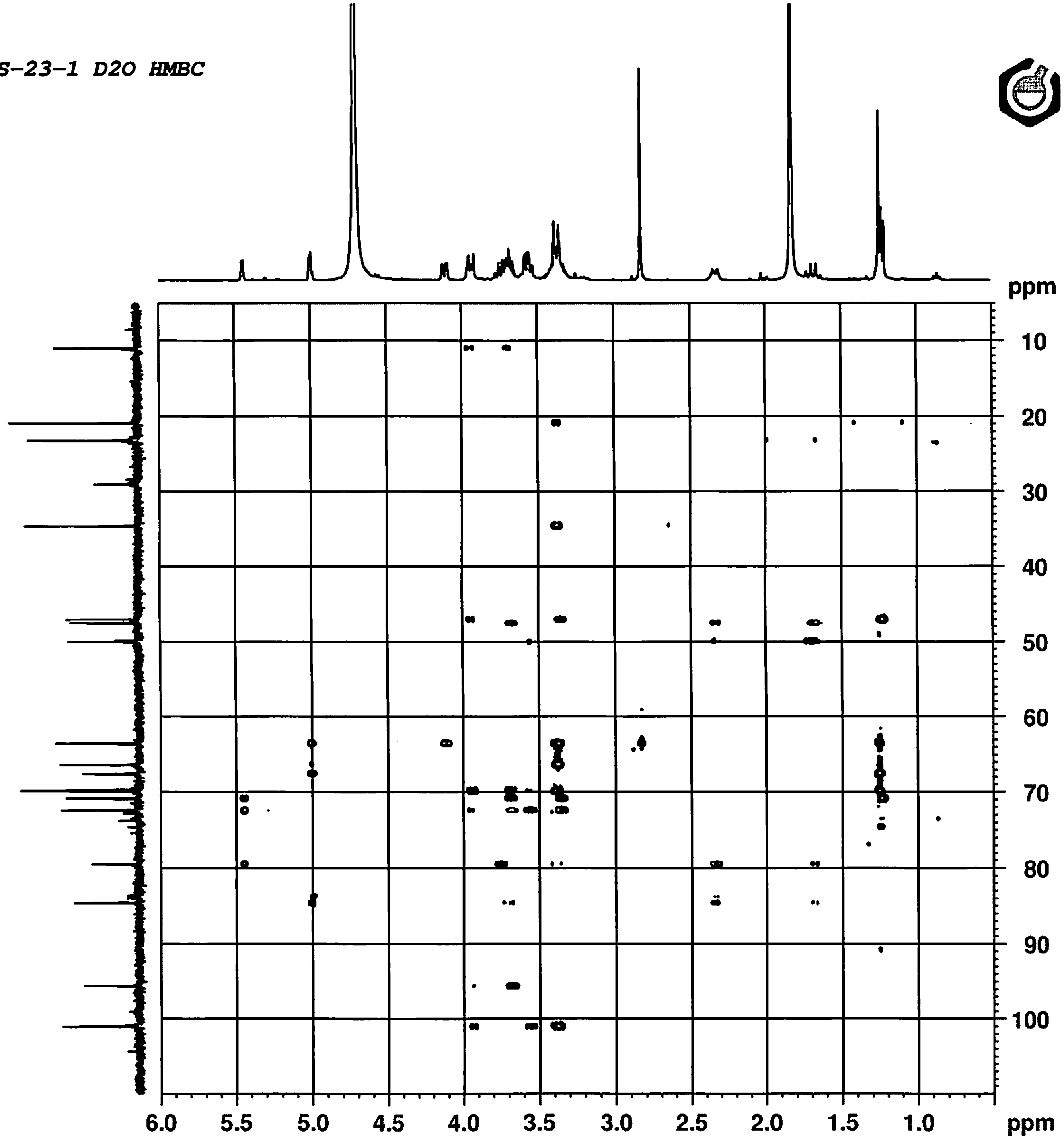


```
Current Data Parameters
NAME       9-MMS-23-1, 20Jun2018
EXPNO                   13
PROCNO                   1

F2 - Acquisition Parameters
Date_             20180621
Time                  2.22 h
INSTRUM              spect
PROBHD    Z116098_0683 (
PULPROG     hmbcetgpl3nd
TD                    4096
SOLVENT                D2O
NS                       8
DS                      16
SWH               6009.615 Hz
FIDRES            2.934382 Hz
AQ               0.3407872 sec
RG                   203.4
DW                  83.200 usec
DE                    6.50 usec
TE                   297.9 K
CNST6          120.0000000
CNST7          170.0000000
CNST13           8.0000000
D0              0.00000300 sec
D1              1.50000000 sec
D6              0.06250000 sec
D16             0.00020000 sec
IN0             0.00002260 sec
TDav                     1
SFO1           400.3326021 MHz
NUC1                    1H
P1                   19.00 usec
P2                   38.00 usec
PLW1            4.09749985 W
SFO2           100.6741315 MHz
NUC2                   13C
P3                   10.00 usec
P24                2000.00 usec
PLW2           66.69200134 W
SPNAM[7]       Crp60comp.4
SPOAL7               0.500
SPOFFS7   0 Hz
SPW7           10.18999958 W
GPNAM[1]        SMSQ10.100
GPZ1                 80.00 %
GPNAM[3]        SMSQ10.100
GPZ3                 14.00 %
GPNAM[4]        SMSQ10.100
GPZ4                 -8.00 %
GPNAM[5]        SMSQ10.100
GPZ5                 -4.00 %
GPNAM[6]        SMSQ10.100
GPZ6                 -2.00 %
P16                1000.00 usec

F1 - Acquisition parameters
TD                     512
SFO1              100.6741 MHz
FIDRES           86.421463 Hz
SW                 219.757 ppm
FnMODE       Echo-Antiecho

F2 - Processing parameters
SI                    2048
SF             400.3300000 MHz
WDW                   SINE
SSB       0
LB        0 Hz
GB        0
PC                    1.40

F1 - Processing parameters
SI                    2048
MC2          echo-antiecho
SF             100.6630586 MHz
WDW                  QSINE
SSB                      2
LB        0 Hz
GB        0
```

Exact mass: 496.27 (736.36 for 4 Acetate salt)

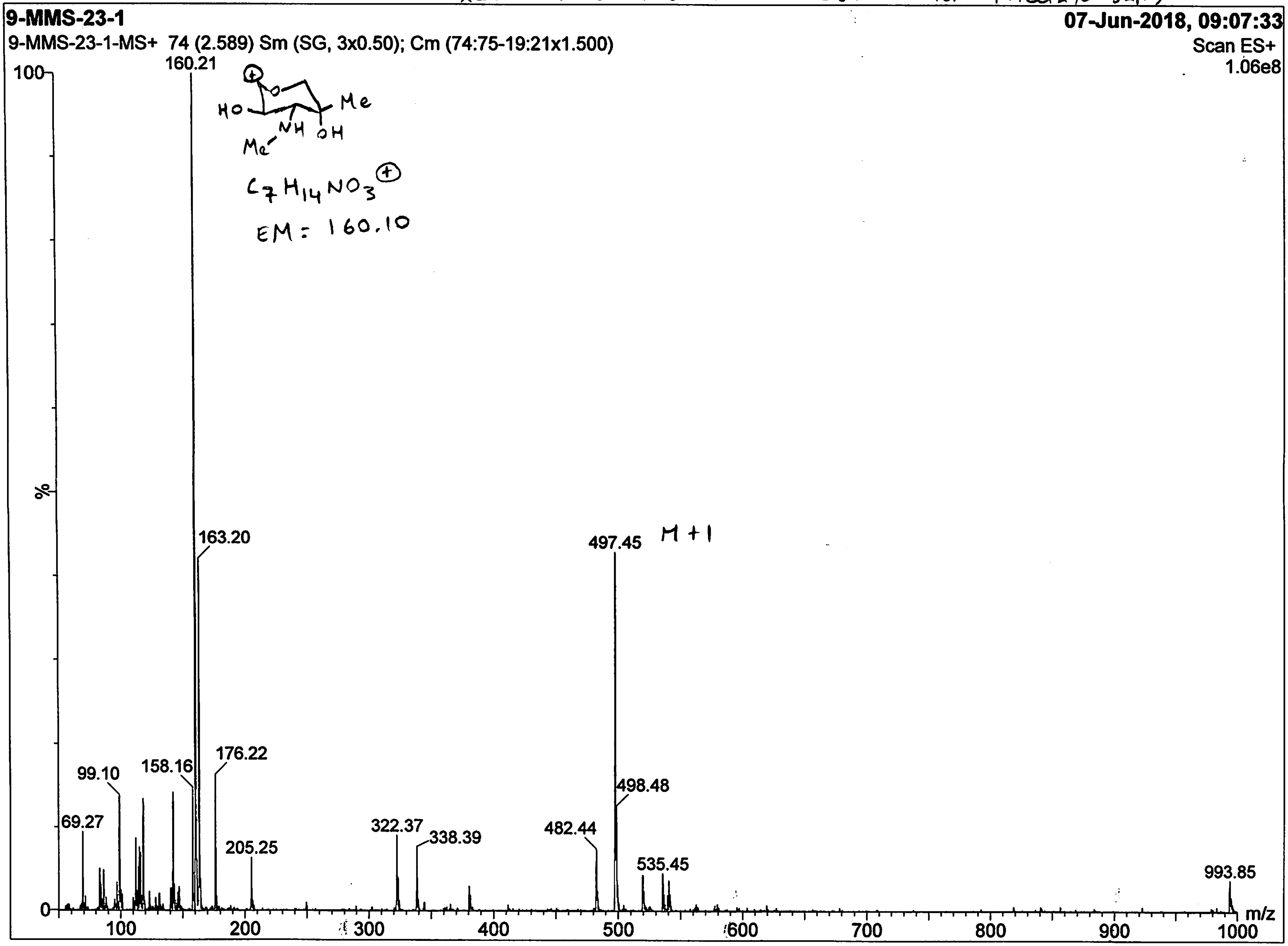

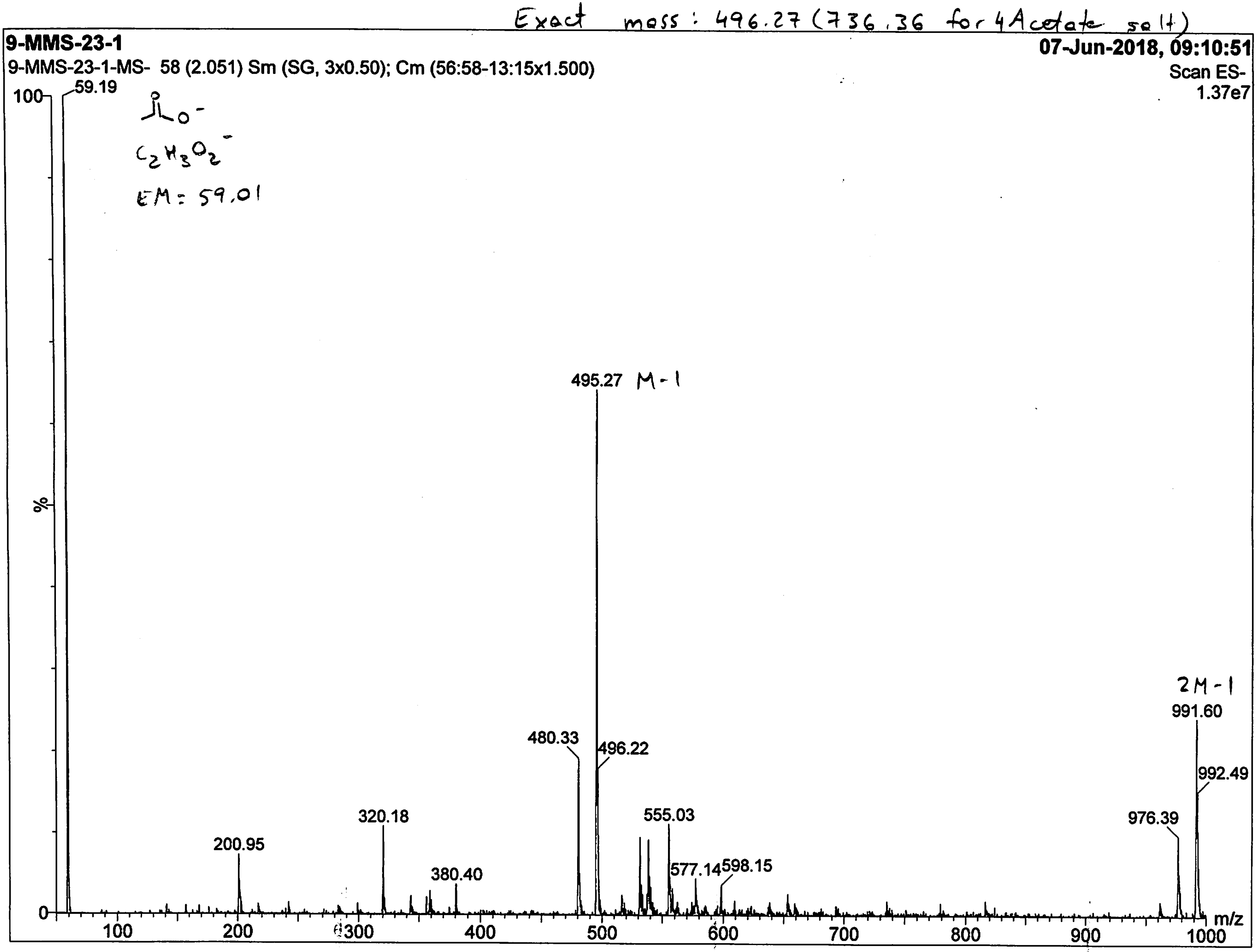
Exact mass: 496.27 (736.36 for 4Acetate salt)
9-MMS-23-1
9-MMS-23-1-MS- 58 (2.051) Sm (SG, 3x0.50); Cm (56:58-13:15x1.500)
07-Jun-2018, 09:10:51
Scan ES-
1.37e7
$C_2H_3O_2^-$
EM = 59.01
59.19
495.27 M-1
480.33
496.22
555.03
577.14
598.15
200.95
320.18
380.40
2M-1
991.60
992.49
976.39
%
m/z
100
200
300
400
500
600
700
800
900
1000
0
100
